# Local cues enable classification of image patches as surfaces, object boundaries, or illumination changes

**DOI:** 10.1101/2025.02.26.640416

**Authors:** Christopher DiMattina, Eden E. Sterk, Madelyn G. Arena, Francesca E. Monteferrante

## Abstract

To correctly parse the visual scene, one must detect edges and determine their underlying cause. Previous work has demonstrated that image-computable neural networks trained to differentiate natural shadow and occlusion edges exhibited sensitivity to boundary sharpness and texture differences. Although these models showed a strong correlation with human performance on an edge classification task, this previous study did not directly investigate whether humans actually make use of boundary sharpness and texture cues when classifying edges as shadows or occlusions. Here we directly investigated this using synthetic image patch stimuli formed by quilting together two different natural textures, allowing us to parametrically manipulate boundary sharpness, texture modulation, and luminance modulation. In a series of initial “training” experiments, observers were trained to correctly identify the cause of natural image patches taken from one of three categories (occlusion, shadow, uniform texture). In a subsequent series of “test” experiments, these same observers then classified 5 sets of synthetic boundary images defined by varying boundary sharpness, luminance modulation, and texture modulation cues using a set of novel parametric stimuli. These three visual cues exhibited strong interactions to determine categorization probabilities. For sharp edges, increasing luminance modulation made it less likely the patch would be classified as a texture and more likely it would be classified as an occlusion, whereas for blurred edges, increasing luminance modulation made it more likely the patch would be classified as a shadow. Boundary sharpness had a profound effect, so that in the presence of luminance modulation increasing sharpness decreased the likelihood of classification as a shadow and increased the likelihood of classification as an occlusion. Texture modulation had little effect on categorization, except in the case of a sharp boundary with zero luminance modulation. Results were consistent across all 5 stimulus sets, showing these effects are not due to the idiosyncrasies of the particular texture pairs. Human performance was found to be well explained by a simple linear multinomial logistic regression model defined on luminance, texture and sharpness cues, with slightly improved performance for a more complicated nonlinear model taking multiplicative parameter combinations into account. Our results demonstrate that human observers make use of the same cues as our previous machine learning models when detecting edges and determining their cause, helping us to better understand the neural and perceptual mechanisms of scene parsing.

## INTRODUCTION

One of the most fundamental tasks of the visual system is to parse the global scene into regions representing distinct surfaces (**Marr, 1982; Frisby & Stone, 2010**). However, at the earlier stages of the visual system, image analysis is very much “local” in nature, with neuronal receptive fields only processing a limited region of the overall scene (**Boussaoud, Desimone, & Ungerleider, 1991; Elston & Rosa, 1998; Kobatake & Tanaka, 1994; Op de Beeck & Vogels, 2000; Wilson & Wilkinson, 2015)**. Nevertheless, local image regions potentially provide critical information for computations essential for global scene segmentation, for instance, edge detection (**Jing et al, 2022; Konishi et al., 2003; Zhou & Mel, 2008; DiMattina, Fox, & Lewicki, 2012; Mely et al., 2016**), region grouping (**Ing, Wilson, & Giesler, 2010**), figure-ground assignment (**Ramenahalli, Mihalas, & Niebur, 2014; Fowlkes, Martin & Malik, 2007; Burge, Fowlkes & Banks, 2010**), and edge classification (**DiMattina et al., 2022; Ehringer et al., 2017; Breuil et al., 2019; Vilankar et al., 2014**). The importance of local image analysis is illustrated in **Fig. 1**, in which we see three image regions whose correct and rapid categorization as an occlusion edge (red square), a cast shadow edge (green square) or a uniformly illuminated surface (blue square) by lower visual areas is potentially useful as input to higher visual areas for segmenting the scene. While a large body of work in vision science and computer vision has focused on identifying locally available cues for edge detection (**Konishi et al., 2003; Martin et al., 2004; Zhou and Mel, 2008**) and figure-ground assignment (**Ramenahalli, Mihalas, & Niebur, 2014; Fowlkes, Martin & Malik, 2007; Burge, Fowlkes & Banks, 2010**), relatively little work has investigated how local edges are classified into different causes like surface boundaries or changes in illumination (**DiMattina et al., 2022; Breuil et al., 2019; Vilankar et al., 2014**). This last problem is essential for accurate scene parsing, as it would be highly detrimental for an organism to mistake a cast shadow for a physical object boundary.

**Figure 1:**
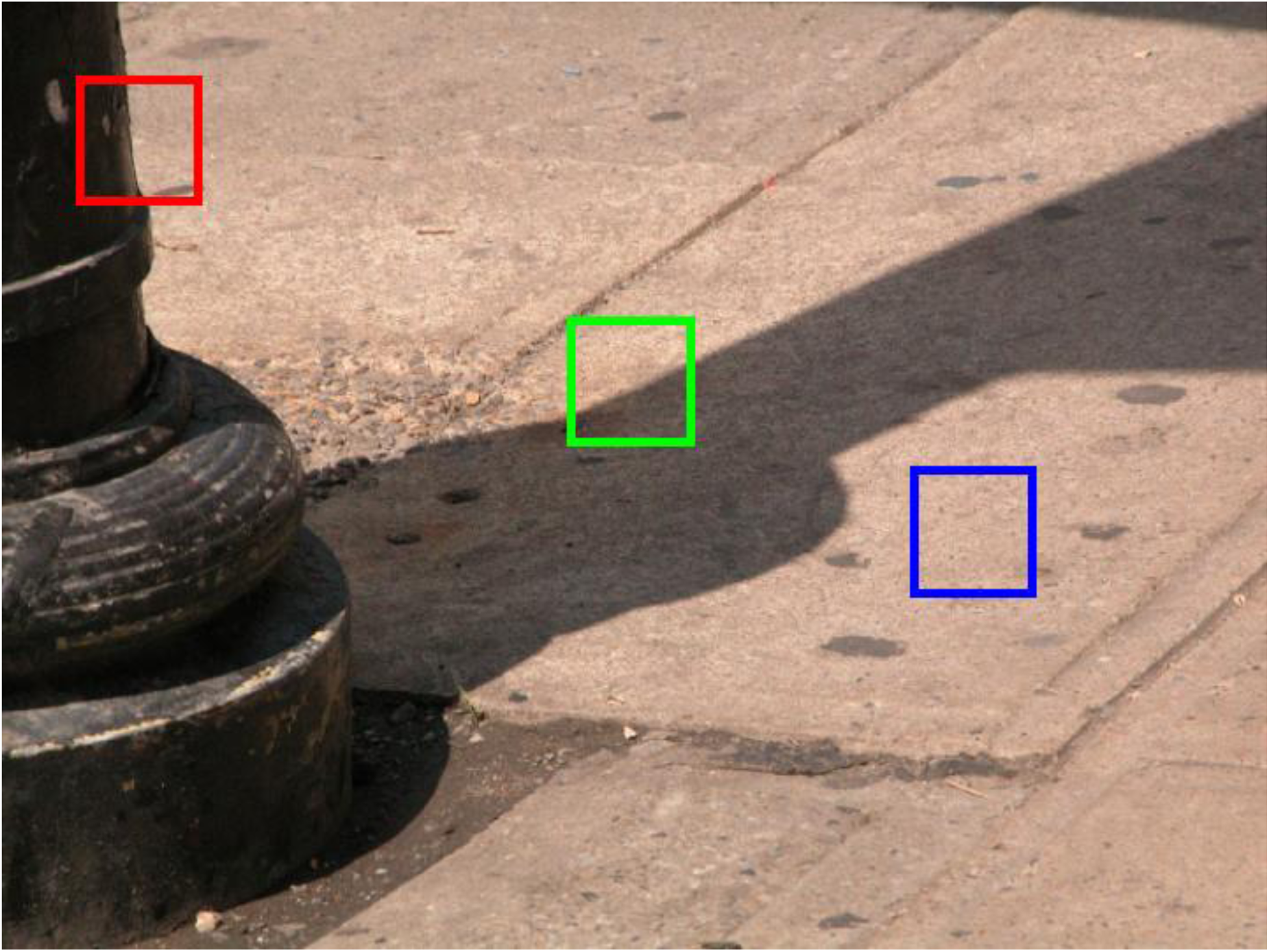
Local image patches arising from three different physical causes. The red square outlines an image patch containing an occlusion boundary (change in material). The green square outlines an image patch containing a cast shadow edge (change in illumination) on a uniform surface. Finally the blue square outlines a singe surface region with no change in illumination.

Although it is well known that changes in color are useful for edge detection (**Zhou & Mel, 2008**; **DiMattina et al., 2012; Hansen & Gegenfurtner, 2009, 2017**), only in the past couple of years has it been clearly demonstrated that changes in color are useful for classification of edges as changes in material or illumination. In a recent study by **Breuil et al. (2019)**, it was shown that performance in classifying edges as illumination or material was greatly enhanced when both color and luminance information were available. This is intuitive, because often at the boundary between two surfaces there are changes in both luminance and chromaticity, whereas a shadow on a uniform surface only gives rise to a change in luminance without an accompanying change in chromaticity. However, color and luminance cues alone are incapable of accounting for human performance in the task, as evidenced by the fact that machine classifiers utilizing these two cues were unable to account for improved human performance observed with increased image patch size (**Breuil et al., 2019**). This is likely because the increased context provided by larger image patches makes other potentially useful cues like texture or penumbral blur more readily perceivable. To better understand the various achromatic cues which may contribute to classifying grayscale edges as occlusions or shadows, **DiMattina et al. (2022)**, measured simple low-level image statistics from 40 × 40 pixel grayscale shadow and occlusion patches, including Michelson contrast, RMS contrast, and proportion of the amplitude spectrum comprised of high-spatial frequencies. Using a simple linear classification method, it was found that these simple statistics can distinguish shadows from occlusions with 70 percent accuracy. These authors also trained image-computable neural network classifiers to distinguish shadows and occlusions, with the first layer weights comprised of multi-scale Gabor filters resembling V1 simple cells. It was found that performance of these image-computable neural network models was in excess of 80 percent, with units in the trained models exhibiting tuning to penumbral blur and texture differences. Comparing model performance to the results of a psychophysical experiment where human observers classified natural image patches as shadows or occlusions, it was found that the models performed as well as the best human observers, with model category probabilities exhibiting significant positive correlations with human category probabilities on an image-by-image basis.

The study by **DiMattina et al. (2022)** established that simple neural network models trained to distinguish shadows and occlusions exhibit tuning to boundary sharpness and texture differences, and their performance correlates well with that of human observers on a natural image patch classification task. However, this work did not establish that humans actually make use of these cues when classifying edges as shadows or occlusions. Therefore, the aim of the current study was to directly test the hypothesis that the cues of luminance, boundary sharpness, and texture differences are utilized by human observers when classifying image patches as shadows, occlusions, or uniformly illuminated textures (surfaces). To do this, we trained human observers on a natural image classification task (with feedback) where they classified natural image patches taken from databases of occlusions, shadows and uniformly illuminated textures (**Olmos & Kingdom, 2004; DiMattina et al., 2012, 2022**). We then required them to classify synthetic image patches (without feedback) generated by quilting together two textures with varying degrees of boundary sharpness, texture modulation, and luminance modulation. This enabled us to directly test the role of each of these parameters (and their various combinations) for local image patch classification. We observed significant effects of each parameter and various combinations of parameters on image patch classification, with image patches having sharp boundaries and strong texture modulation being likely to be classified as occlusions, image patches with blurred boundaries and intermediate levels of luminance modulation being likely to be classified as shadows, and patches without texture or luminance modulation being classified as uniform surfaces. Fitting multinomial logistic regression models to our behavioral data revealed significant effects of these variables, and these models were able to generalize well to predict the categorization of stimuli not used for model training. This work demonstrates that humans are making use of the same locally available cues of luminance modulation, boundary sharpness, and texture modulation, extending the results from our previous study (**DiMattina et al., 2022**) and confirming our hypothesis about the central importance of these achromatic cues.

## METHODS

### Visual stimuli

#### Natural images

Previously developed databases of occlusions, shadows and textures (**Olmos & Kingdom, 2004; DiMattina et al., 2012; DiMattina et al., 2022**) were used to obtain 40 × 40 image patches from each of these categories. These image patches were used to define two training surveys which participants completed in order to learn the classification task (additional details in the next section). A set of N = 100 images (768 × 576 pixels) containing occlusions and uniformly illuminated textures (surfaces) taken from **DiMattina et al. (2012)** and a set of N = 45 images (768 × 576 pixels) containing shadows (**DiMattina et al., 2022**) was used in the current study. Training Surveys 1 and 2 (**TRN-1**, **TRN-2**) were constructed by sampling 50 image patches (40 × 40 pixels) containing each kind of stimulus (occlusion, shadow, texture) from the images in our set, so that each survey contained 150 image patches. Examples of images patches from **TRN-1** from each category are shown in **Fig. 2**, and patches from **TRN-2** are shown in **Supplementary Fig. S1**.

**Figure 2:**
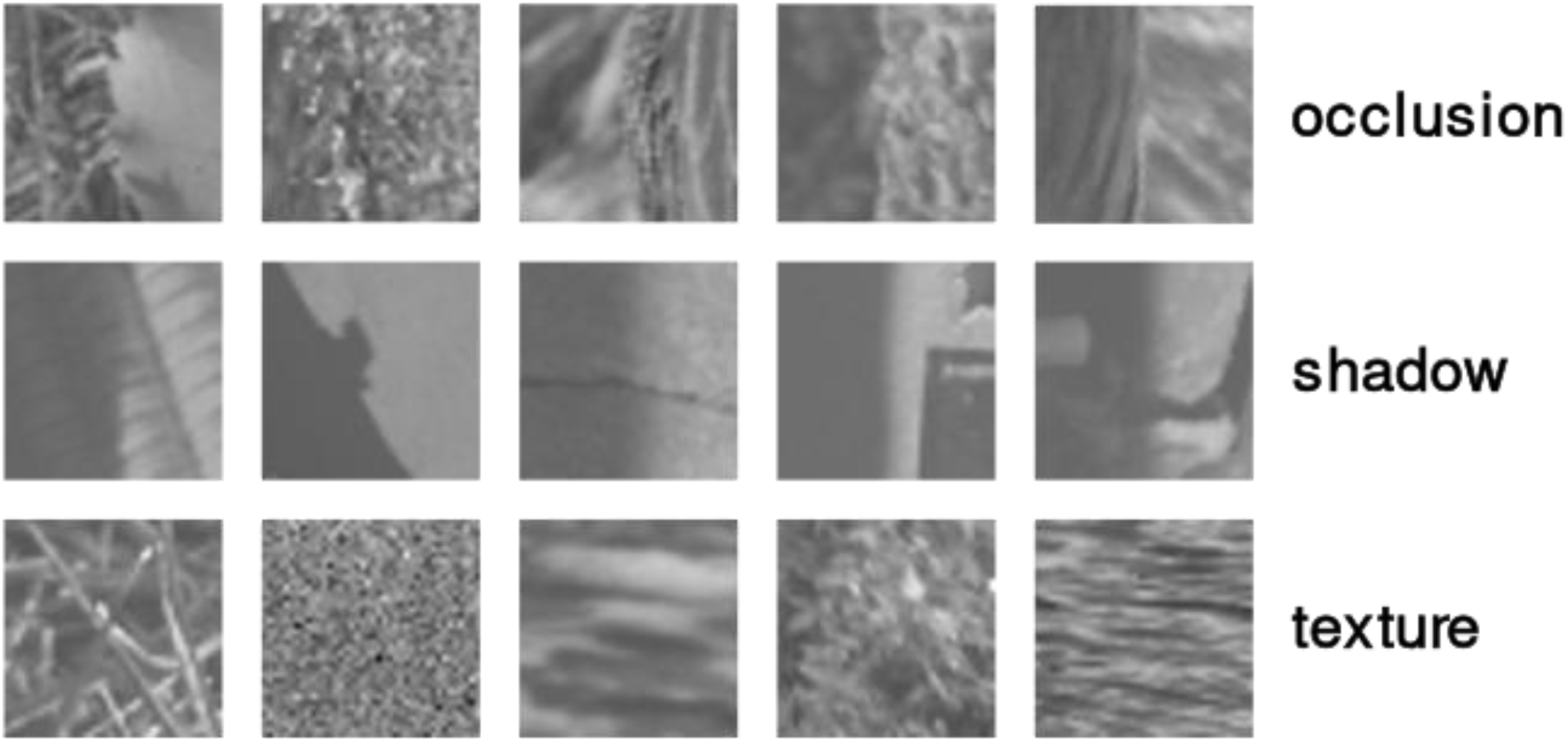
Examples of stimuli from Training Survey 1 (**TRN-1**) from each of the three categories.

#### Synthetic images

We created synthetic image patches by quilting together two different 40 × 40 texture patches from the McGill database (**Olmos & Kingdom, 2004**) on opposite sides of a vertical boundary. These stimuli allowed us to parametrically vary the degree of texture and luminance differences on opposite sides of the boundary, as well as the sharpness of the boundary. Depending on the values of these parameters, individual image patches could resemble occlusions, shadows, or uniformly illuminated textures (which we will also refer to as *surfaces*). These synthetic image patches were used to create stimuli for 5 test surveys which we used to study the effects of various image manipulations on classification performance (additional details in next section). A subset of the images from Test Survey 1 (**TST-1**) are shown in **Fig. 3**. Mathematically, given textures *T*_1_and *T*_2_ (each with zero mean luminance and 20 percent RMS contrast), a new texture *T*_*Q*_ was defined in column *j* by

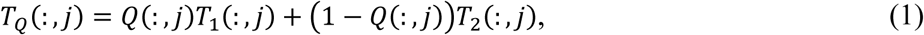

**Figure 3:**
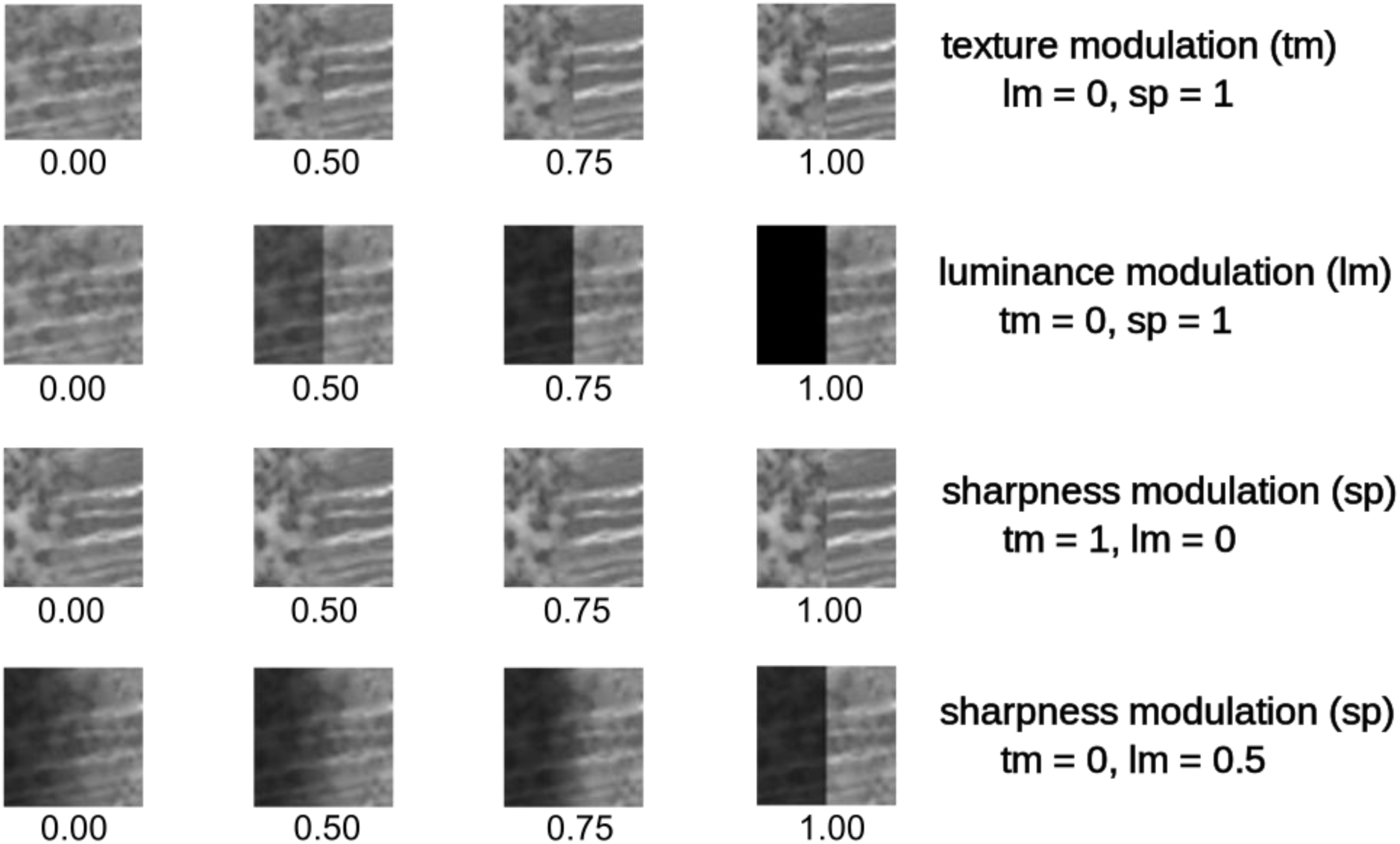
Synthetic image patches from Test Survey 1 (**TST-1**) which illustrate the effects of varying specific parameters. Top row: Effects of varying the degree of texture modulation (**tm**) with no luminance modulation (**lm** = 0) at maximum sharpness (**sp** = 1). Second row: Effects of varying luminance modulation (**lm**) with no texture modulation (**tm** = 0) and maximum sharpness (**sp** = 1). Third row: Effects of varying sharpness (**sp**) in a boundary with maximum texture modulation (**tm** = 1) and no luminance modualtion (**lm** = 0). Fourth row: Effects of varying sharpness (**sp**) in patch with no texture modulation (**tm** = 0) and maximum luminance modulation (**lm** = 1).

where *Q*(:, *j*)is an envelope defined by the equations

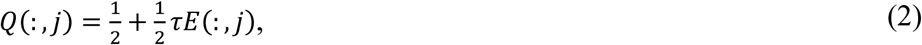

where

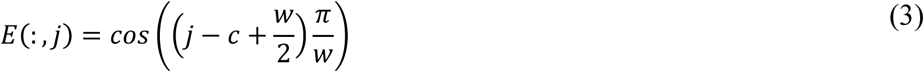

for *c* − *w*⁄2 ⩽ *j* ⩽ *c* + *w*⁄2, with *E*(:, *j*) = 1for *j* ⩽ *c* − *w*⁄2and *E*(:, *j*) = −1for *j* ⩾ *c* + *w*⁄2. The parameter 0 ≤ τ ≤ 1 in **Eq. 2** determines the depth of the texture modulation (**tm**). When τ = 0, there is no texture modulation and the quilted texture *T*_*Q*_ is simply the arithmetic mean of *T*_1_ and *T*_2_. When τ = 1, there is complete texture modulation so that on the left, the quilted patch is identical to *T*_1_ and on the right side is identical to *T*_2_. In **Eq. 3**, *c* is the center pixel and *w* denotes the width of the taper window in pixels, which is determined by *w* = (1 − σ)*N*, where *N* is the number of pixels in the image. Here the parameter 0 ≤ σ ≤ 1 determines the sharpness (**sp**), or 1 minus the proportion of the image over which the two textures transition into each other. σ = 1 corresponds to an abrupt transition between textures so that the left side is *T*_1_, the right side is *T*_2_, and there is no central region over which they are mixed. σ = 0 corresponds to a gradual mixing of the two textures on the left and right over the entire extent of the patch.

Finally, we manipulate the luminance modulation (**lm**) on opposite sides of the image patch. This is accomplished by multiplying the quilted texture *T*_*Q*_ by a luminance modulation envelope L, defined by

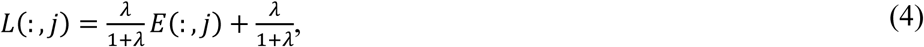

so that our final stimulus image I is given by the element-wise product (⊙) of L and *T*_*Q*_:

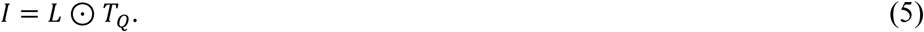

In **Eq. 4**, the parameter 0 ≤ λ ≤ 1 determines the extent of luminance modulation. λ = 0 corresponds to no luminance modulation, whereas λ = 1 corresponds to maximum luminance modulation where one side of the image patch is black and the other has maximum luminance. Therefore, the three parameters τ, σ, and λ allow one to manipulate the texture modulation, boundary sharpness, and luminance modulation and hence vary the appearance of the texture patch so that it resembles a shadow, occlusion, or uniform surface, as we can see in **Fig. 3**.

### Participants and Experimental Design

#### Participants

Participants consisted of N = 67 students at Florida Gulf Coast University enrolled in undergraduate Psychology classes during the Spring 2024 semester. These participants completed online surveys using Qualtrics (https://www.qualtrics.com) in which they had to classify image patches as being occlusions, shadows, or textures (see below for additional details). All participants provided informed consent prior to each survey, and all procedures were approved beforehand by the FGCU Institutional Review Board (IRB Protocol 2021-78), in accordance with the Declaration of Helsinki.

#### Surveys

To familiarize participants with the three image categories, participants first took two training surveys in which they had to classify 40 × 40 grayscale natural image patches as being occlusions, shadows, or textures. Occlusion, texture, and shadow patches were taken from hand-labeled databases utilized in previous research (**DiMattina, 2012**; **DiMattina et al., 2022**). Each training survey (**Training Survey 1**, **Training Survey 2**) was comprised of 150 questions (50 occlusions, 50 shadows, 50 textures), and participants were given feedback after each question indicating the correct categorization of each image patch. The goal of providing feedback was for each participant to learn how natural occlusions, shadows, and textures can appear, and what cues define these natural categories. Examples of stimuli from each of these categories from Training Survey 1 (**TRN-1**) are shown in **Fig. 2**. Examples image patches from Training Survey 2 (**TRN-2**) are shown in **Supplementary Fig. S1**. A sample question from **TRN-1** is shown in **Supplementary Fig. S2**. After completing both training surveys, participants completed 5 test surveys. In each test survey, participants had to classify 40 × 40 gray-scale computer-generated image patches as being occlusions, shadows, or textures. Upon classifying each patch, participants were not given feedback on how they answered. Each test survey (**TST-1**,…,**TST-5**), was comprised of 64 questions, each containing a single image patch. A subset of the images in the Test Survey 1 (**TST-1**) is shown in **Fig. 3**, and a sample question is shown in **Supplementary Fig. S3**. Only those participants who had successfully completed both training surveys were included in our analysis of the test survey results shown here.

### Logistic regression models

We sought to build a simple predictive model which would relate the defining parameters of the synthetic image patches to the probability of that patch being classified by human observers as each of the three categories (occlusion, shadow, texture). Perhaps the simplest such model is the multinomial logistic regression (MLR) model, which generalizes binomial logistic regression (**Bishop, 2006; Murphy, 2012**). In the most basic application of this model to our problem, there are *N* = 3 predictor variables **lm** (*u*_1_), **sp** (*u*_2_), **tm** (*u*_3_) and *K* = 3 possible categorization behaviors (occlusion, shadow, texture). The log-probability of classification in category *k* = 1, 2, 3 is given by a linear combination of the predictor variables

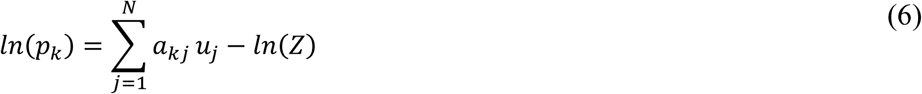

where *a*_*kj*_ are the linear coefficients, and *Z* is a normalization constant, given by

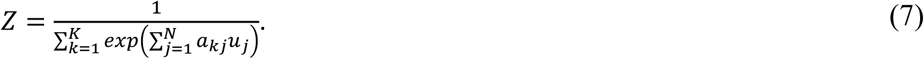

The machine learning problem is to use categorization probabilities *p*_*k*_ for each of the synthetic image patches along with the values of the parameters (*u*_1_, *u*_2_, *u*_3_) used to define the synthetic image patch to find the linear coefficients *a*_*kj*_ that minimize the difference between the observed and predicted log-probabilities. Both models were fit to our psychophysical data making use of methods implemented in R (https://www.r-project.org/). The fitting methods implemented in R simplify the machine learning problem by fitting the log-odds ratio of two of the three categories relative to the reference category. In our analysis, the reference category was occlusions (*k* = 1). Therefore, the fitting the model defined in **Eq. 6, 7** reduces fitting a set of two linear equations with 8 free parameters, defined in **Eq. 8, 9**.

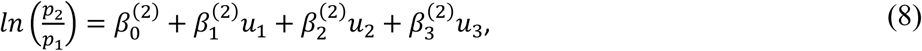

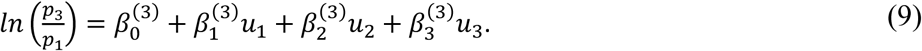

In order to determine how well the model defined in **Eq. 8** and **Eq. 9** could predict observer responses for datasets not used for model fitting, we formed 5 Jackknife Sets (**JCK-1**,…,**JCK-5**) by pooling all of the test surveys, excluding one test survey (for instance, **JCK-1** pools **TST-2** to **TST-5**, excluding **TST-1**). The model was fit to each of the 5 Jackknife sets (**JCK-i**), and used to predict observer responses for the excluded test survey (**TST-i**).

We also fit a more complicated multinomial logistic regression model to our data. This model was similar to the one in **Eq. 8**, **9**, but also included all possible two-way and three-way interactions between variables, as defined in **Eq. 10, 11**.

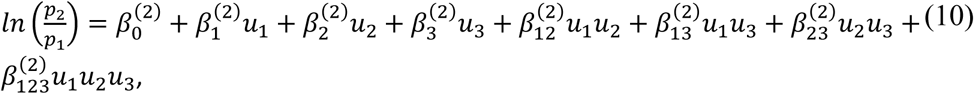

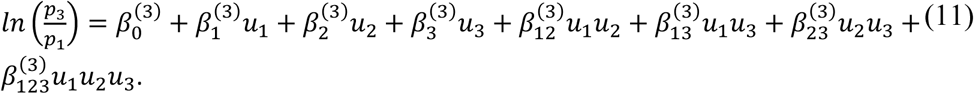

We refer to the logistic regression model defined by **Eq. 8, 9** as the *linear model*, since the predictor variables are simply those used to define the image patches, and the model defined by **Eq. 10, 11** as the *nonlinear model*, since the predictor variables include all possible products of the variables used to define the image patches. It is important to note however that both models are linear in the coefficients, and hence can be estimated using standard methods. In order to assess whether there was any improvement in model fitting, while penalizing for the increased model complexity of the nonlinear model, we compared the two models using the AIC (Akaike Information Criterion), a measure which rewards good fits to data while penalizing for model complexity (**Akaike, 1974**; **Bishop, 2006)**. The formula for the AIC is given by the expression

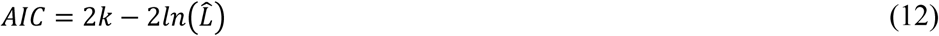

Where *k* is the number of model free parameters, and *L*^^^ is the value of the data likelihood function at its maximum value. As we see from **Eq. 12**, a smaller value of the AIC corresponds to a better complexity-penalized fit.

## RESULTS

### Classification of natural image patches

N = 66 participants completed Training Survey 1 (**TRN-1**) and N = 66 completed Training Survey 2 (**TRN-2**), with N = 65 completing both surveys. **Fig. 4** shows for each of the three types of image patches (occlusions, shadows, textures) the probability of it being classified as such by the participants. The leftmost panel of **Fig. 4a** shows the probability that image patches of each kind (occlusions = red, shadows = green, textures = blue) in **TRN-1** are classified as occlusions, sorted by the probability of being classified as such. Similarly, the middle and rightmost panels of **Fig. 4** show the probability that each of the image patches in **TRN-1** are classified as shadows (middle) or textures (right), sorted by the probability of being classified as such. **Fig. 4b** is the same as **Fig. 4a** but for **TRN-2**.

**Figure 4:**
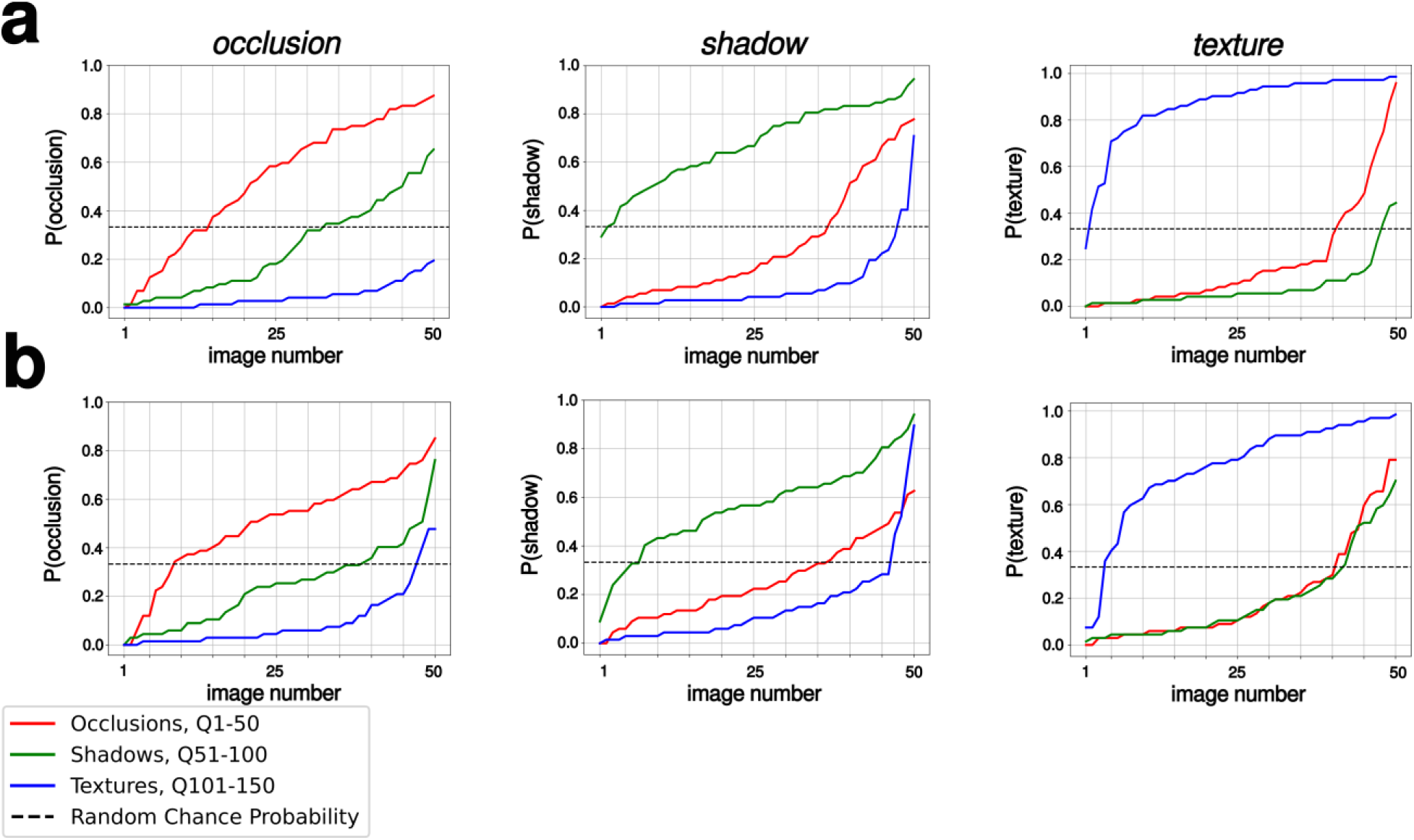
Proportions that the image patches of each kind in the Training sets (red: occlusions, green: shadows, blue: textures) were classified as each of the three categories (left: occlusions, center: shadows, right: textures). Image patches from each category were sorted in order of their probability of being classified as the column category. (**a**) Training Survey 1 (**TRN-1**) (**b**) Training Survey 2 (**TRN-2**).

**Table 1** and **Table 2** show the confusion matrices for the classification task for training surveys **TRN-1** and **TRN-2**, respectively. From both **Fig. 4** and these tables that on the whole, participants perform well above chance in classifying the 40 × 40 image patches into three categories, with 69.79% percent correct classifications for **TRN-1**, and 61.48% for **TRN-2**. For **TRN-1**, occlusions were correctly classified 54.78% of the time, but were misclassified as shadows 26.39% of trials and as textures 18.81% of trials. Similarly, for **TRN-2**, we find occlusions to be correctly classified 51.30% of the time, but were misclassified as shadows 25.72% of trials and textures 22.96% of trials. Therefore, the most common mistake for occlusions was to misclassify them as shadows, although a substantial number were misclassified as textures. We see from **Table 1** and **Table 2** that shadows were somewhat more likely to be correctly classified (**TRN-1**: 67.6%, **TRN-2**: 56.21%), with the more common mistake to be to misidentify them as occlusions (**TRN-1**: 24.6%, **TRN-2**: 23.48%) rather than misidentifying them as textures (**TRN-1**: 7.78%, **TRN-2**: 20.3%). Finally, textures were most likely to be correctly identified as such (**TRN-1**: 87.0%, **TRN-2**: 76.93%), with relatively few misclassified as occlusions (**TRN-1**: 4.48%, **TRN-2**: 9.03%) or shadows (**TRN-1**: 8.51%, **TRN-2**: 14.03%).

**Table 1:** Confusion matrix for Training Survey 1 (TRN-1). Actual categories are in rows, classified categories are in columns.

|  | Occlusion | Shadow | Texture | Total |
| --- | --- | --- | --- | --- |
| Occlusion | 1808 | 871 | 621 | 3300 |
| Shadow | 812 | 2231 | 257 | 3300 |
| Texture | 148 | 281 | 2871 | 3300 |

**Table 2:** Confusion matrix for Training Survey 2 (TRN-2). Actual categories are in rows, classified categories are in columns.

|  | Occlusion | Shadow | Texture | Total |
| --- | --- | --- | --- | --- |
| Occlusion | 1693 | 849 | 758 | 3300 |
| Shadow | 775 | 1855 | 670 | 3300 |
| Texture | 298 | 463 | 2539 | 3300 |

### Classification of synthetic image patches

#### Effects of luminance cues

The effects of each of the three cues on the probability that a synthetic image patch was classified as an occlusion, shadow, or texture were highly dependent on the values of the other cues. **Fig. 5** shows the effect of luminance modulation for three values (0, 0.5, 1) of texture modulation (*rows*) and sharpness (*columns*), averaged over all 5 test surveys. Remarkably consistent results were obtained for all of the individual surveys (**Supplementary Figs. S4-S6**), justifying our averaging procedure. As we see in the leftmost column of **Fig. 5**, with or without texture modulation, when the boundary between regions was not sharp (**sp** = 0.0), as luminance was increased we observed a decrease in the likelihood of classification as uniform texture (blue curve), and increase in the likelihood of classification as a shadow (green curve). However, as we see in the upper right panel, for a sharp boundary with no texture modulation, as we increase the luminance cue the image patch is less likely to be classified as a uniform surface and more likely to be classified as an occlusion. Finally, we see that when there is a sharp boundary with strong texture modulation, it is almost always classified as an occlusion, independent of the luminance modulation value (bottom row, rightmost column).

**Figure 5:**
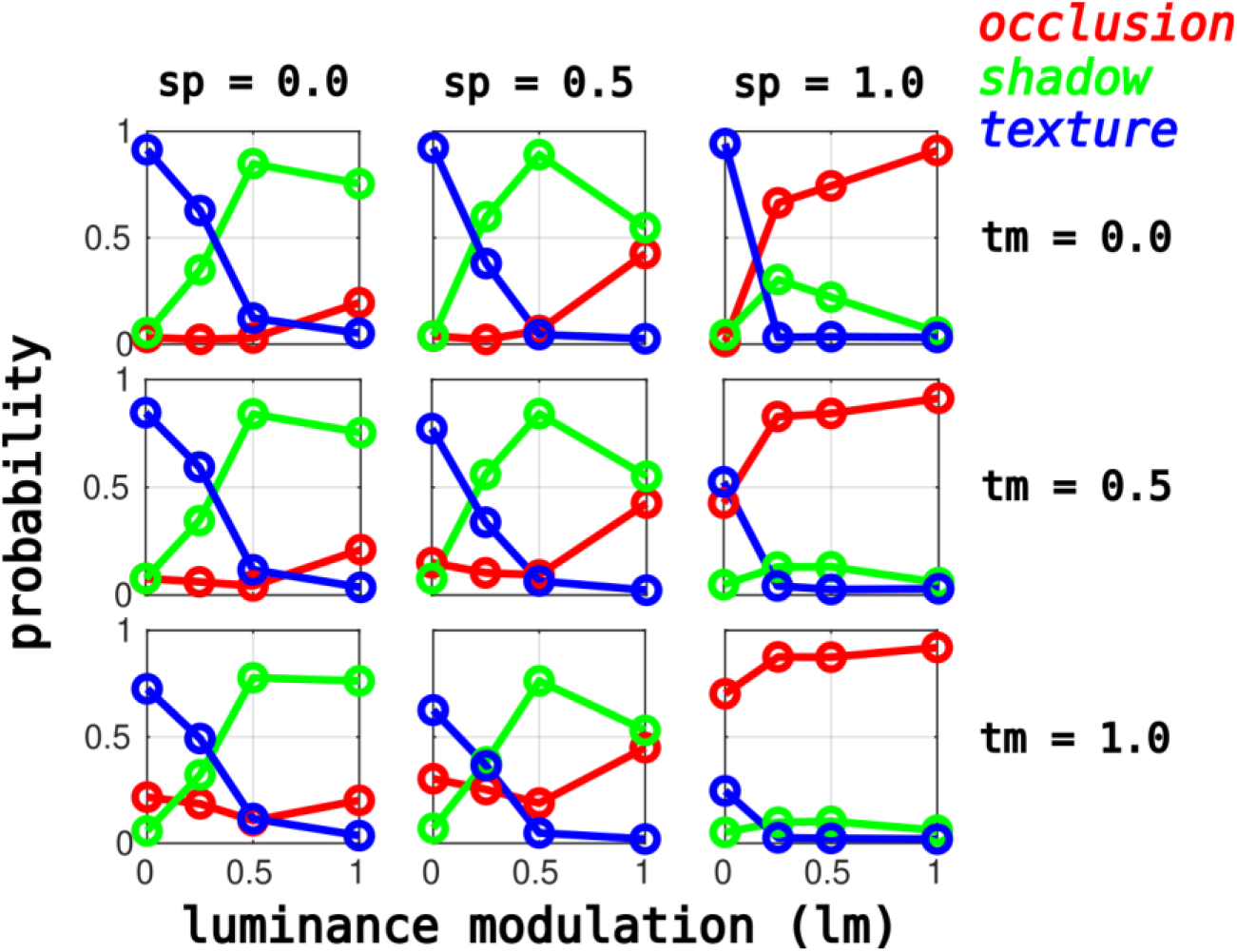
Interaction of variables for image patch categorization probability, averaged across all 5 test surveys. Plots show probability of an image patch being classified as an occlusion (red lines), shadow (green lines), or surface (blue lines) as a function of luminance modulation (**lm**) for the endpoint (0, 1) and midpoint (0.5) values of sharpness (**sp:** columns) and texture modulation (**tm:** rows).

We can also visualize the effects of luminance from an investigation of **Fig. 8**, where the normalized (R, G, B) values of the colored squares surrounding each image patch are proportional to the probabilities of the image patch in **TST-1** being classified as an occlusion, shadow, and surface, respectively. We see that in the absence of luminance cues (**lm** = 1.0, upper left), as one increases boundary sharpness the classification interpolates between surface (blue) to occlusion (red), except in the case where there is no texture modulation (**tm** = 1.0). In cases with non-zero luminance modulation (upper right, bottom), we see that regardless of texture modulation, as boundary sharpness is increased the image patch goes from being classified as a shadow (green) to an occlusion (red). Similar results were obtained for **TST-2** to **TST-5** (**Supplementary Fig. S7-S10**).

#### Effects of texture cues

As we see from **Fig. 6**, texture modulation (**tm**) has little effect on categorization probability, except in the absence of luminance modulation (top row). In this case, we see for sharp boundaries (**sp** = 1.0) lacking luminance cues (**lm** = 0.0), as one increases texture modulation one reduces the probability of classification as a texture (blue curve) and increases the probability of classification as an occlusion (red curve). This can also be seen by examining **Fig. 9**. For luminance modulation greater than zero, we see that for all non-zero levels of texture modulation (upper right, bottom) as one increases boundary sharpness the classification interpolates between shadows (green) and occlusions (red). For zero luminance modulation (**lm** = 0.0), in the absence of texture modulation, the image patch will be classified as a surface (**tm** = 0.0, upper left panel), but in the presence of texture modulation, as the boundary becomes sharper the patch goes from being classified as a surface to being classified as an occlusion (upper right panel, bottom). Similar results were obtained for **TST-2** to **TST-5** (**Supplementary Fig. S11-S14**).

**Figure 6:**
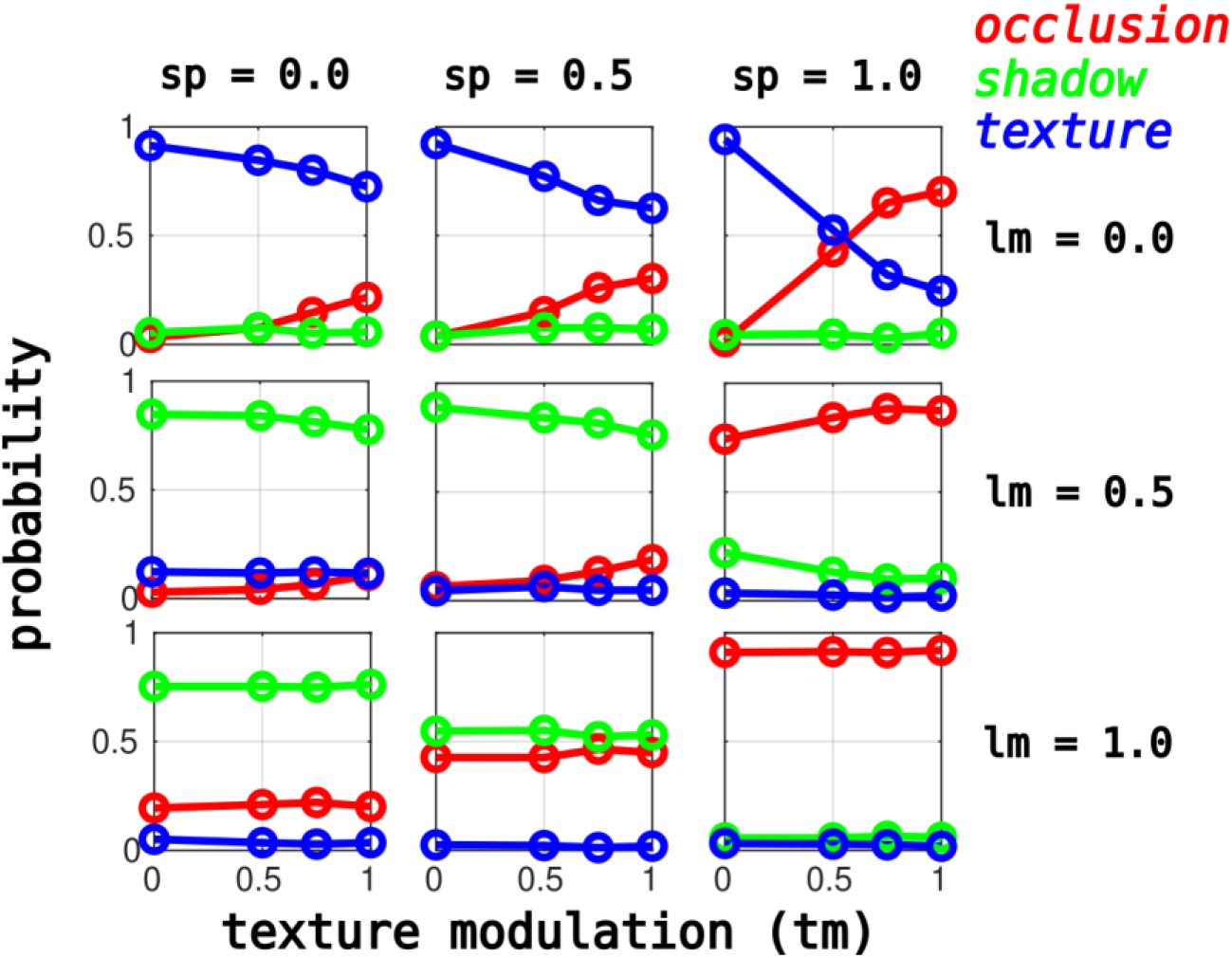
Interaction of variables for image patch categorization probability, averaged across all 5 test surveys. Plots show probability of an image patch being classified as an occlusion (red lines), shadow (green lines), or surface (blue lines) as a function of texture modulation (**tm**) for the endpoint and midpoint values of sharpness (**sp:** columns) and luminance modulation (**lm:** rows).

#### Effects of boundary sharpness

Boundary sharpness has a profound effect on categorization probability, although its effect depends on the luminance modulation level. As we see in **Fig. 7** (top row), in the absence of luminance modulation, for high levels of texture modulation (**tm** = 1.0), as one increases the boundary sharpness, the image patch is more likely to be interpreted as an occlusion (red curves) than a texture (blue curve). In the presence of luminance modulation, then regardless of the level of texture modulation, we see that as the boundary sharpness varies from 0.0 to 1.0, the patch goes from being interpreted as a shadow (green curves) to being interpreted as an occlusion (red curves). One can also see that **Fig. 10** that for maximally sharp boundaries (**sp** = 1.0), in the presence of any texture or luminance modulation, the image patch is very likely to be classified as an occlusion (red, bottom right). We also see that for less sharp boundaries, regardless of texture modulation, as luminance is increased the image patch goes from being classified as a surface (blue) to a shadow edge (green). Similar results were obtained for **TST-2** to **TST-5** (**Supplementary Fig. S15-S18**).

**Figure 7:**
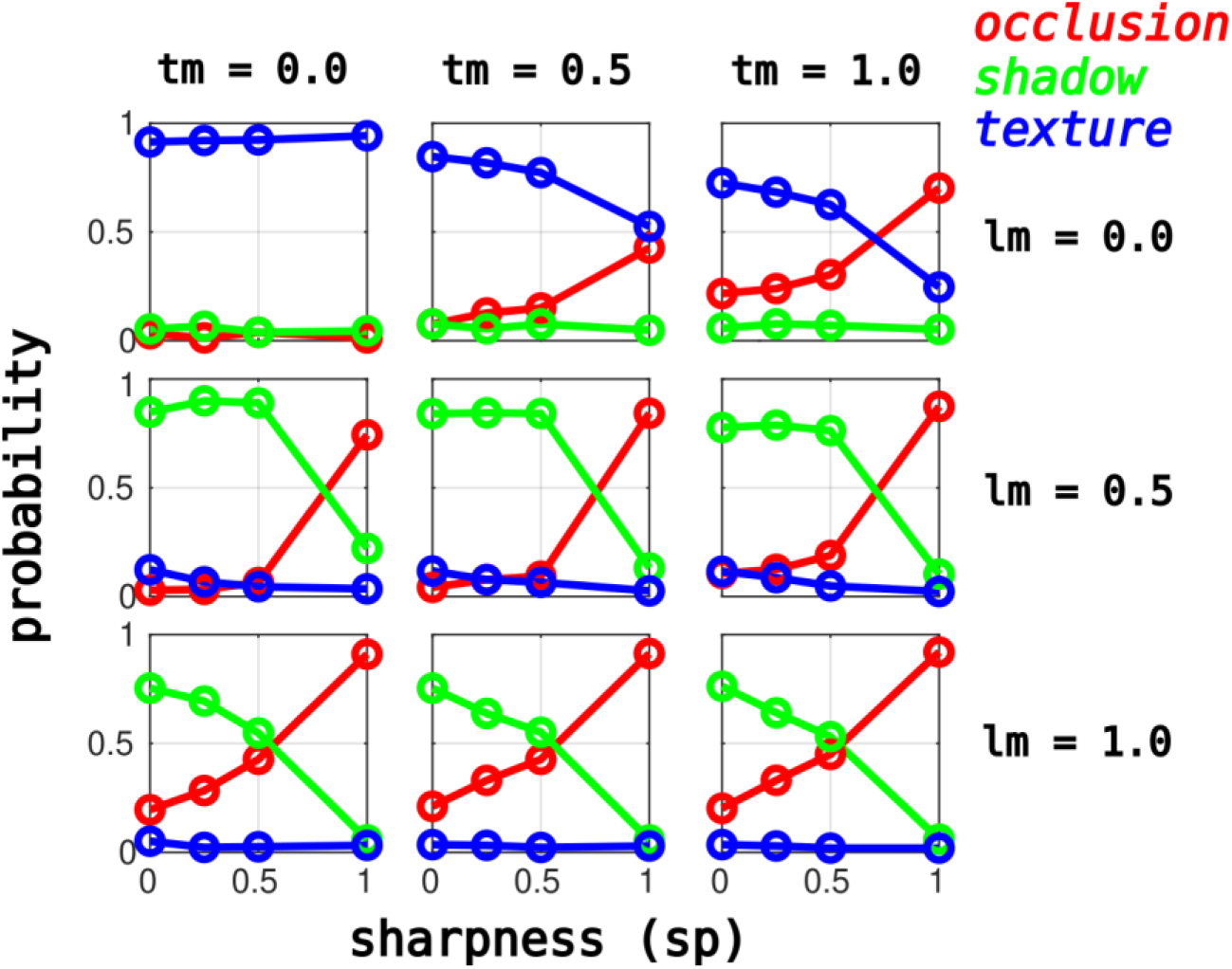
Interaction of variables for image patch categorization probability, averaged across all 5 test surveys. Plots show probability of an image patch being classified as an occlusion (red lines), shadow (green lines), or surface (blue lines) as a function of sharpness (**sp**) for the endpoint and midpoint values of texture modulation (**tm:** columns) and luminance modulation (**lm:** rows).

**Figure 8:**
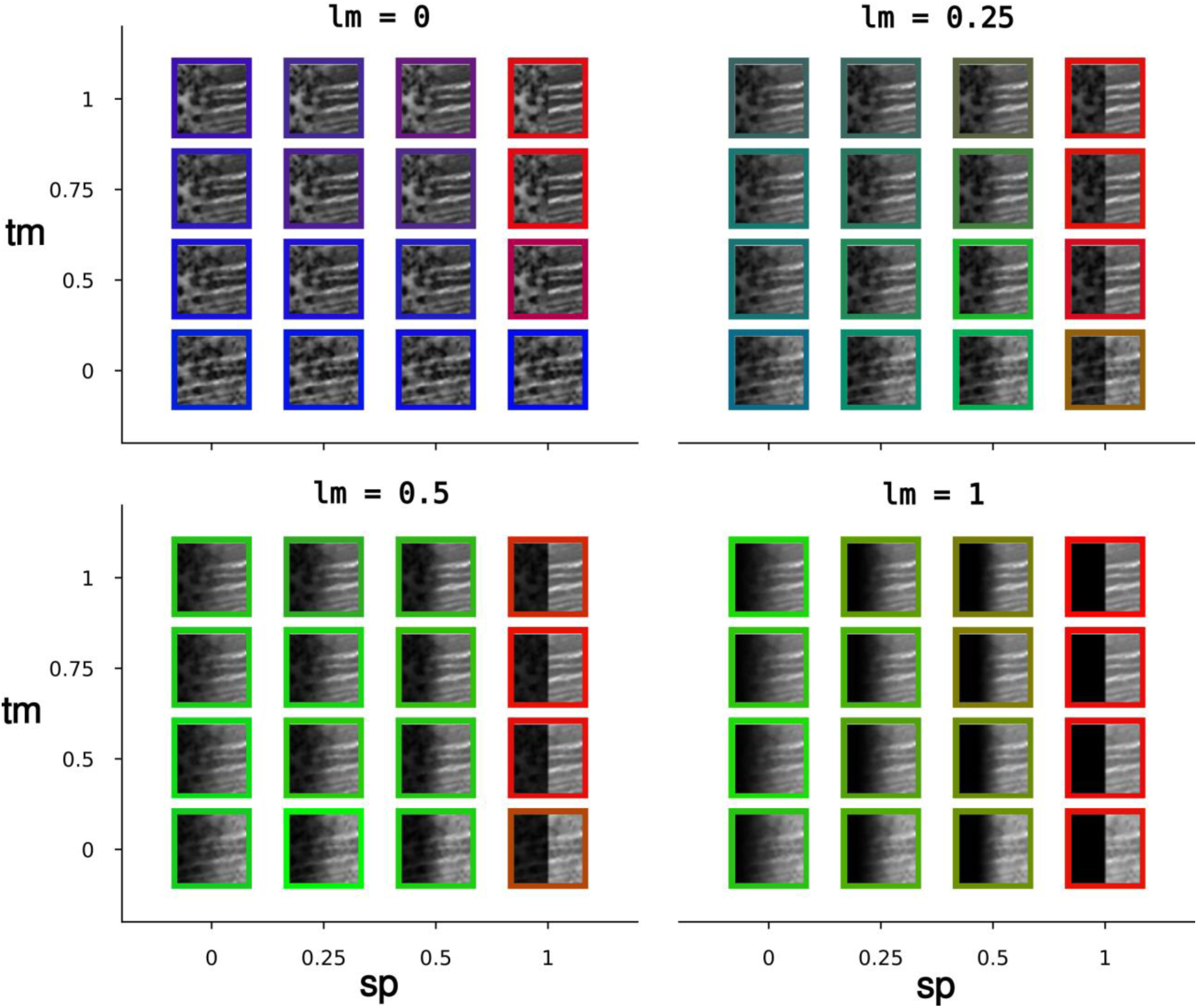
All of the image patches from **TST-1**, together with the values of their defining parameters. The (R,G,B) color of the border is determined by the proportion of times that the patch was classified as an occlusion (R-value), shadow (G-value) or surface (B-value). Here the image patches are grouped by the level of the luminance modulation (**lm**).

**Figure 9:**
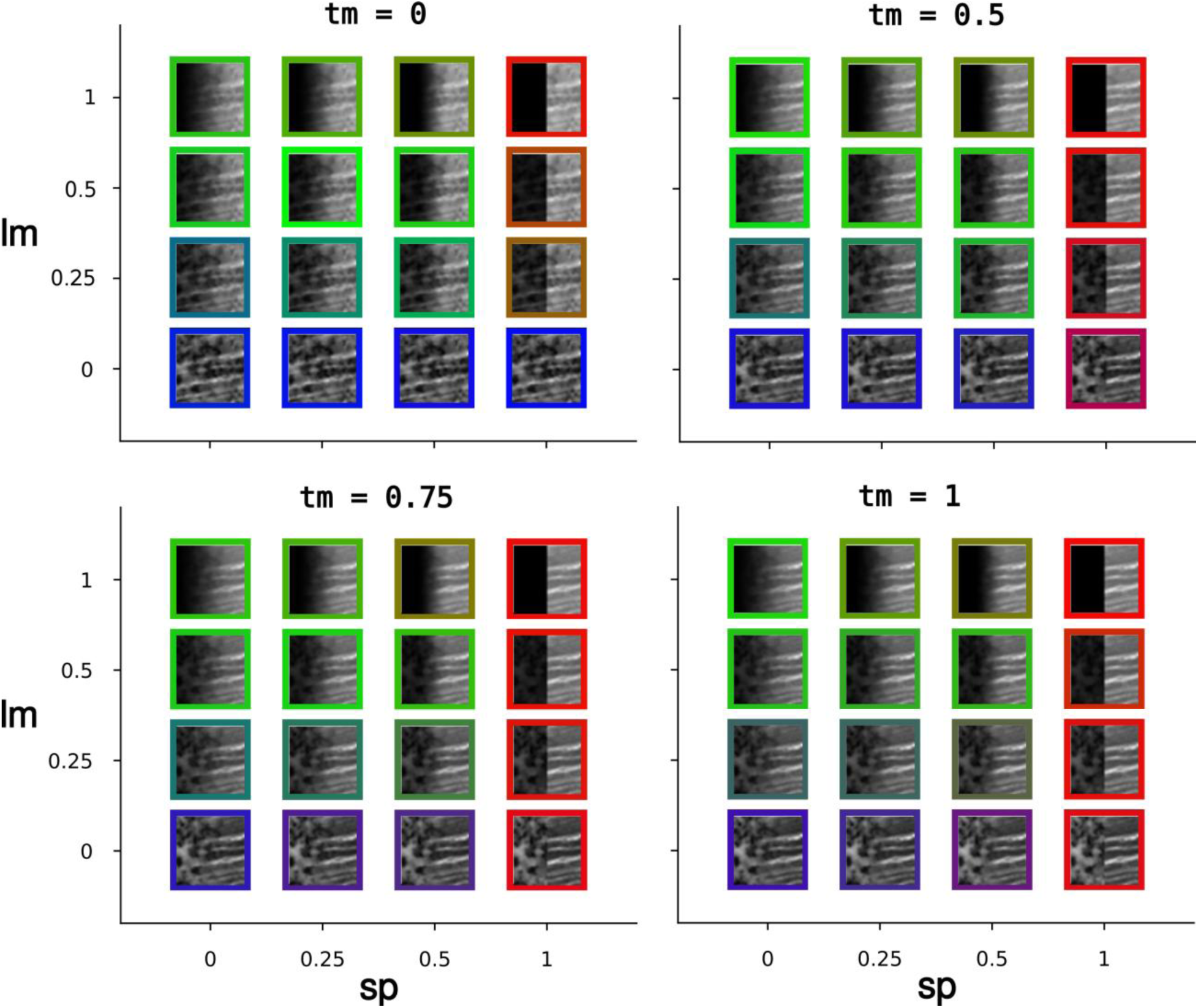
All of the image patches from **TST-1**, together with the values of their defining parameters. The (R,G,B) color of the border is determined by the proportion of times that the patch was classified as an occlusion (R-value), shadow (G-value) or surface (B-value). Here the image patches are grouped by the level of the texture modulation (**tm**).

**Figure 10:**
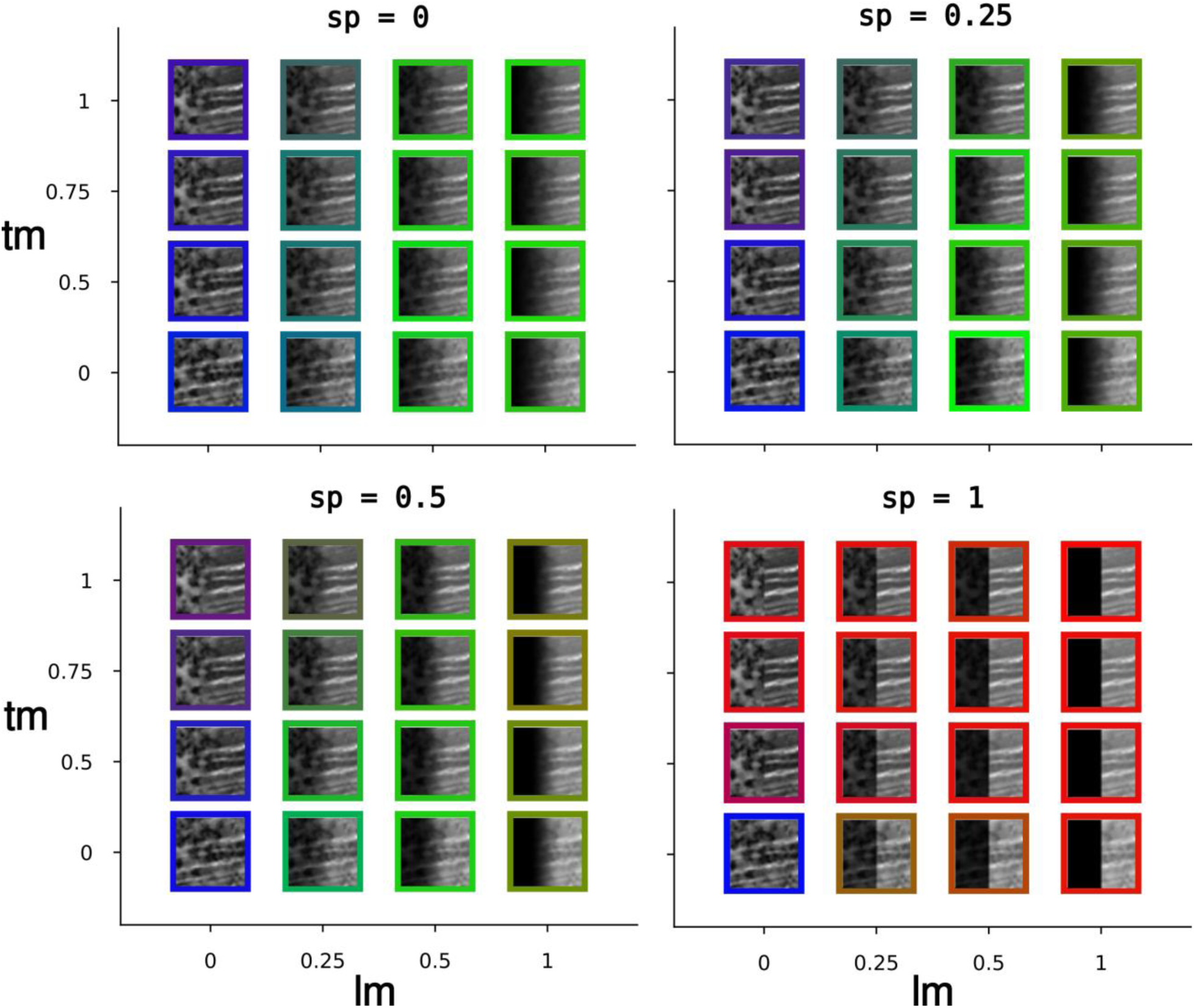
All of the image patches from **TST-1**, together with the values of their defining parameters. The (R,G,B) color of the border is determined by the proportion of times that the patch was classified as an occlusion (R-value), shadow (G-value) or surface (B-value). Here the image patches are grouped by the level of the boundary sharpness.

### Multinomial linear regression models

We next tried to determine whether our classification results could be accounted for by some model defined on the three variables defining the image patches. Perhaps the simplest such model is a linear multinomial logistic regression model (with 8 free parameters) defined on the variables defining the image patches (**lm**, **tm**, **sp**), as described in **Eq. 8** and **Eq. 9**. A more complex nonlinear model (16 free parameters) was also fit to the data (**Eq. 10, 11**), with this model including all possible combinations of products of the defining variables. For each of the five test surveys, the models were fit to all of the other surveys excluding one which was held back (**Jackknife Sets 1-5**), and then the fitted model was used to predict the results of the hold-out test survey.

We found that qualitatively both the basic and derived models do a good job of predicting the results of surveys not used to train the model. The implementation of multinomial logistic regression in R fits the log of the ratio of each category (shadow, texture) relative to the reference category (occlusion). **Table 3** shows the coefficients of the linear model fit to **Jackknife Set 1 (JCK-1)**. We see from these coefficients in the first row that increasing the luminance modulation and texture modulation slightly decreases the likelihood of being classified as a shadow relative to an occlusion, whereas increasing the sharpness of the boundary greatly decreases the likelihood of a shadow relative to an occlusion. This is consistent with our observations in **Figs. 5-7** which show that increasing boundary sharpness strongly favors classification as an occlusion rather than a shadow. We also see from the coefficients in the second row that increasing luminance modulation and boundary sharpness drastically decrease the likelihood of being classified as a shadow relative to an occlusion, again consistent with our results in **Figs. 5-7**. Coefficients for fits to all other Jackknife Training Sets are shown in **Tables 4-7**, and we observe similar results to those in **Table 3**. **Fig. 11** - **Fig. 13** show observer classifications (solid lines, circles) and model predictions (dashed lines, diamonds) for **TST-1**, where the model was fit to **JCK-1**. We see a strong agreement between model predictions and observer performance.

**Figure 11:**
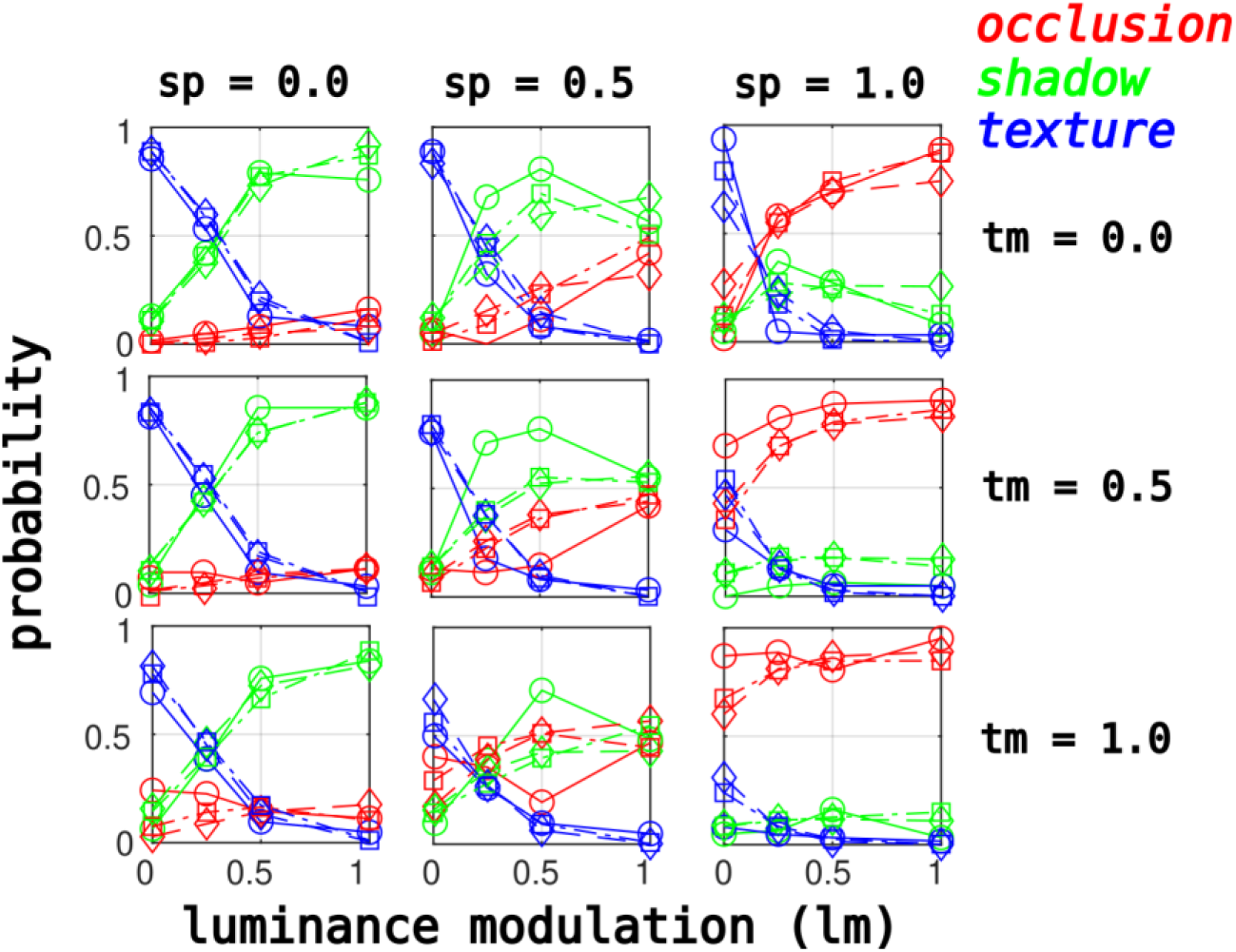
Observer performance on **TST-1** (circles, solid lines), together with the predictions of a linear multinomial logistic regression model (diamonds, dashed lines) and nonlinear multinomial logistic regression models (squares, dash-dot lines). Plot shows luminance modulation on the horizontal axis.

**Figure 12:**
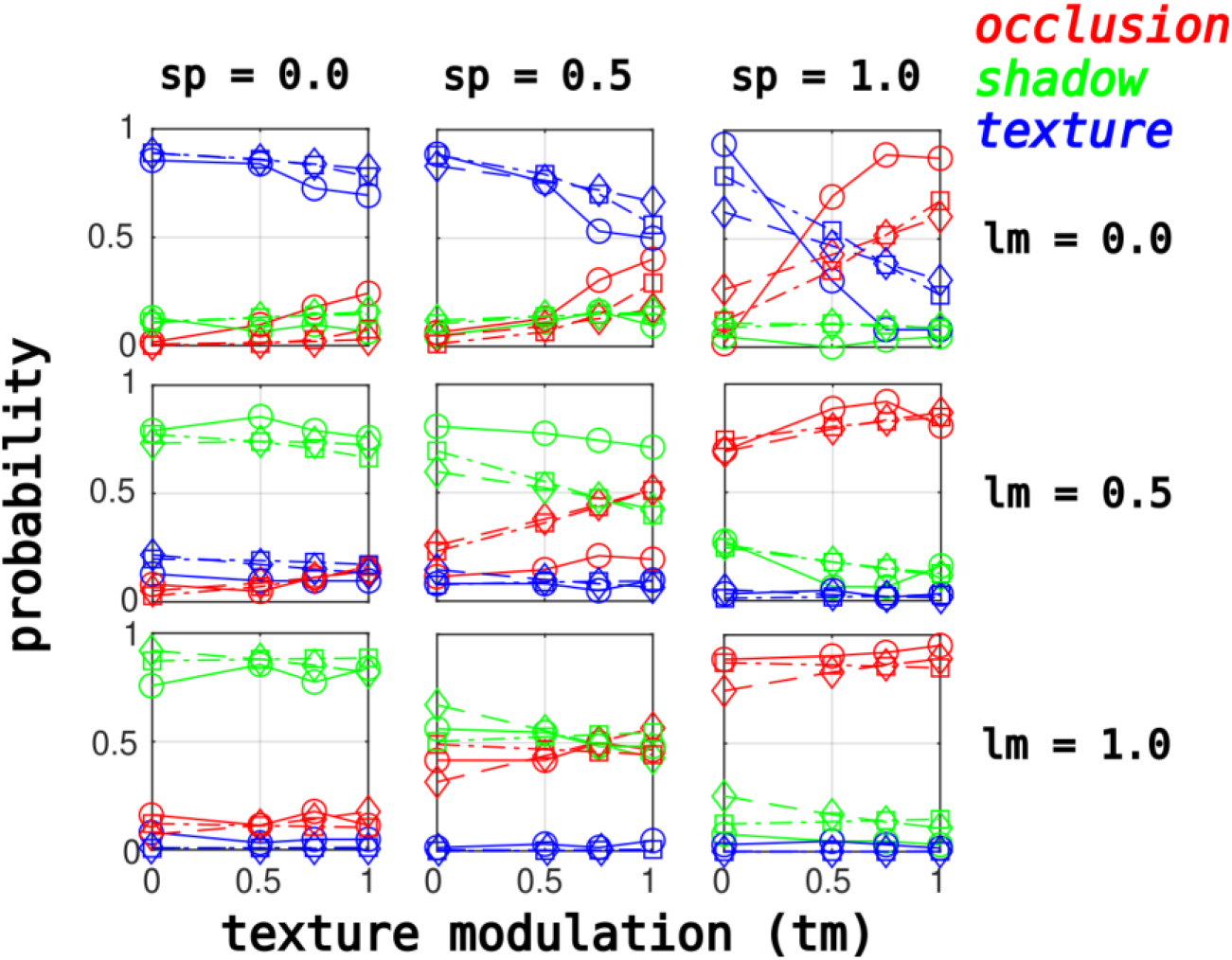
Observer performance on **TST-1** (circles, solid lines), together with the predictions of a linear multinomial logistic regression model (diamonds, dashed lines) and nonlinear multinomial logistic regression models (squares, dash-dot lines). Plot shows texture modulation on the horizontal axis.

**Figure 13:**
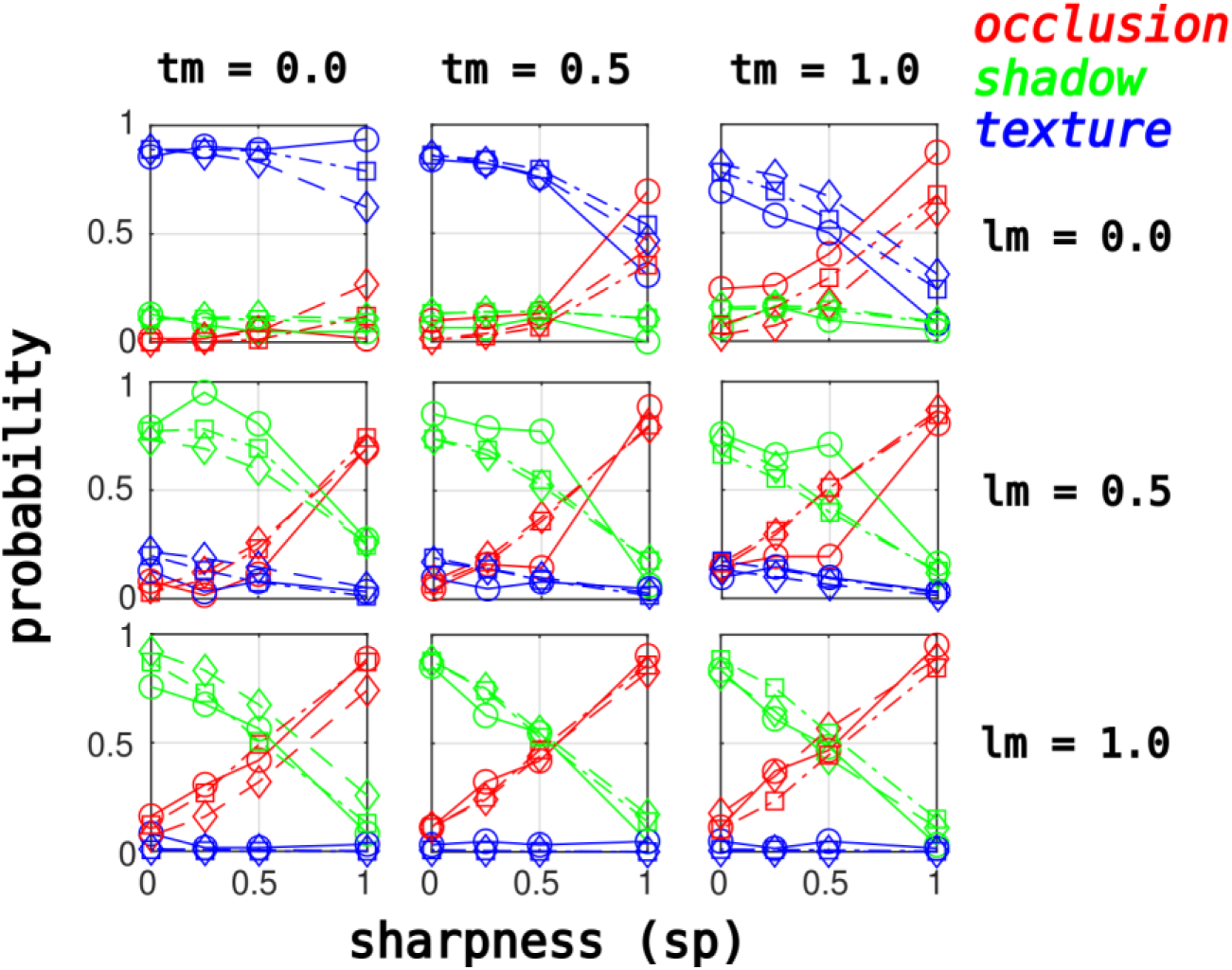
Observer performance on **TST-1** (circles, solid lines), together with the predictions of a linear multinomial logistic regression model (diamonds, dashed lines) and nonlinear multinomial logistic regression models (squares, dash-dot lines). Plot shows sharpness on the horizontal axis.

**Table 3:** Coefficients (+/- standard error) for basic linear model for the log-odds ratio of two categories (shadow, texture) relative to the reference category (occlusion) for model fitted to Jackknife Set 1 (JCK-1). A negative coefficient means that increasing the value of the variable decreases the likelihood of that category relative to the likelihood of an occlusion, whereas a positive coefficient increases the likelihood of that category relative to an occlusion.

|  | **Intercept** | **lm** | **sp** | **tm** |
| --- | --- | --- | --- | --- |
| **Shadow** | 2.739 +/- 0.075 | -0.178 +/- 0.066 | -3.626 +/- 0.068 | -1.028 +/- 0.063 |
| **Texture** | 4.875 +/- 0.087 | -6.879 +/- 0.130 | -4.026 +/- 0.083 | -1.516 +/- 0.075 |

**Table 4:** Same as Table 3, but for JCK-2.

|  | **Intercept** | **lm** | **sp** | **tm** |
| --- | --- | --- | --- | --- |
| **Shadow** | 2.681 +/- 0.072 | 0.002 +/- 0.064* | -3.638 +/- 0.067 | -1.139 +/- 0.062 |
| **Texture** | 4.567 +/- 0.084 | -6.435 +/- 0.127 | -3.872 +/- 0.082 | -1.647 +/- 0.075 |

\* = $p > 0.05$ (fails to reach significance)

**Table 5:** Same as Table 3, but for JCK-3 and TST-3.

|  | **Intercept** | **lm** | **sp** | **tm** |
| --- | --- | --- | --- | --- |
| **Shadow** | 2.562 +/- 0.073 | 0.098 +/- 0.065* | -3.619 +/- 0.068 | -1.130 +/- 0.063 |
| **Texture** | 4.541 +/- 0.084 | -6.114 +/- 0.122 | -3.915 +/- 0.082 | -1.590 +/- 0.074 |

\* = $p > 0.05$ (fails to reach significance)

**Table 6:** Same as Table 3, but for JCK-4 and TST-4.

|  | **Intercept** | **lm** | **sp** | **tm** |
| --- | --- | --- | --- | --- |
| **Shadow** | 3.190 +/- 0.080 | -0.579 +/- 0.069 | -3.968 +/- 0.072 | -0.922 +/- 0.065 |
| **Texture** | 5.340 +/- 0.093 | -7.294 +/- 0.133 | -4.480 +/- 0.088 | -1.342 +/- 0.077 |

**Table 7:** Same as Table 3, but for JCK-5 and TST-5.

|  | **Intercept** | **lm** | **sp** | **tm** |
| --- | --- | --- | --- | --- |
| **Shadow** | 2.582 +/- 0.071 | 0.006 +/- 0.063* | -3.490 +/- 0.066 | -1.171 +/- 0.062 |
| **Texture** | 4.431 +/- 0.082 | -6.038 +/- 0.121 | -3.706 +/- 0.080 | -1.762 +/- 0.074 |

\* = $p > 0.05$ (fails to reach significance)

In addition to the simple model defined by **Eq. 8** and **Eq. 9**, we also fit a more complex nonlinear model which regresses not only on the values of the variables, but also on the values of pairwise products and three-way products, as defined in **Eq. 10**, **Eq. 11**. This more complex model better fit the data than the simpler model, as evidenced by the values of the AIC for each model shown in **Table 8**, with smaller AIC values indicating a better fit when a penalty is applied for model complexity. However, as we can see from **Fig. 11** - **Fig. 13**, both the linear (dashed lines, diamonds) and nonlinear models (dashed-dot lines, squares) seem to fit reasonably well. Analogous results are shown in **Supplementary Figs S19-S22** for all other test surveys, with luminance modulation on the horizontal axis.

**Table 8:** AIC from fits of the linear and nonlinear models to each of the 5 Jackknife sets.

| **Jackknife Set** | **Linear Model** | **Nonlinear Model** |
| --- | --- | --- |
| **1** | 23289.28 | 22900.81 |
| **2** | 23901.89 | 23450.94 |
| **3** | 23497.64 | 23026.66 |
| **4** | 22482.04 | 22159.84 |
| **5** | 24114.99 | 23639.25 |

## DISCUSSION

### Overview and Summary

Complex natural stimuli are defined by multiple features, and therefore it is often unclear which features are most useful for distinguishing between two or more natural stimulus categories. One useful approach for gaining traction on such questions is to create synthetic naturalistic stimuli whose various features can be manipulated independently in order to determine which features are most important for classification tasks. This approach has been widely used to better understand the neural and behavioral representations of complex natural stimuli like human speech (**Lieberman, 1996**), animal vocalizations (**Margoliash & Fortune, 1992; DiMattina & Wang, 2006; Osmanski & Wang, 2023**), and human faces (**Loffler et al., 2005; Peterson et al., 2022**). In the current study, we developed a set of novel naturalistic image patch stimuli whose defining features could be modified systematically, enabling us to investigate the role of multiple cues for classifying image patches as being object boundaries, shadow edges, or uniformly illuminated surfaces. To the best of our knowledge, this approach is novel, as previous psychophysical studies addressing these issues have relied on natural image patch token stimuli whose features cannot be varied parametrically (**DiMattina et al., 2012; Vilankar et al., 2014; Breuil et al., 2019; DiMattina et al., 2022**).

A number of previous studies have compared the statistical properties of different edge categories (**Vilankar et al., 2014; Ehinger et al., 2017; Breuil et al., 2019; DiMattina et al., 2022**), or compared the statistical differences between edges and surfaces (**DiMattina et al., 2012; Mely et al., 2016**). Recently, **DiMattina et al. (2022)** addressed the somewhat narrower problem of what cues distinguish edges arising from changes in illumination (cast shadows) versus those arising from object boundaries (occlusions). A statistical analysis of differences between shadow and occlusion edges revealed different distributions of luminance, and differences in the spatial frequency content. Training image-computable neural network models to distinguish shadow patches and occlusion patches revealed that the units in the network were sensitive to boundary sharpness and texture differences. Furthermore, these relatively simple network models (2 or 3 layers deep) were able to perform the classification task as well as the best humans, and exhibited strong positive correlations with human classification decisions on an image-by-image basis. However, this study did not directly examine whether or not the cues which the networks are sensitive to (texture, boundary sharpness) were cues which human participants were using to classify images as shadows or occlusions. Therefore, in the present study, we utilized parametric stimuli in order to directly demonstrate that all three of these cues are utilized by human observers for classifying relatively small (40 × 40) image patches.

We found that luminance modulation was important for distinguishing uniformly illuminated surfaces from shadow edges and occlusion boundaries, although it did not strongly distinguish occlusions from shadows, as evidenced by the relatively small (and sometimes insignificant) coefficients obtained from fitting the linear model to the 5 Jackknife training sets (**Tables 3-7**). Boundary sharpness was found to reliably distinguish occlusions from shadows, as well as distinguishing occlusions from uniform textures in the presence of texture modulation. This is sensible since often the physics of shadow formation gives rise to a blurry boundary between regions of high and low illumination (**Elder & Zucker, 1998**). Finally, texture modulation, although less useful than the other two cues for category classification (as evidenced by smaller model coefficients in **Tables 3-7**), still played a useful role in enabling one to distinguish uniform surfaces from occlusions – even with a sharp boundary, in the absence of texture modulation, one will classify a stimulus as a surface, whereas the presence of texture modulation enables its classification as a boundary. This is consistent with our analysis in **DiMattina et al. (2022)**.

### Limitations and Future Directions

One limitation that is inherent to any study making use of artificial naturalistic stimuli is that they are not exact replicas of natural stimuli, and it is possible that there may be aspects of natural occlusions or shadows that are not well captured by the stimuli we used. However, for several reasons, we think that our stimuli do a reasonably good job of emulating the natural categories. One reason is that as we see from **Figs. 5-7**, for many values of the parameters (usually the endpoints), there is almost universal agreement as to what category the stimulus is. Furthermore, there is a remarkable degree of consistency between different test surveys (**Supplementary Fig. S4-S6**), despite each survey being comprised of different pairs of textures. Finally, as we see in **Fig. 3**, the stimuli certainly do bear a strong resemblance to all three categories, depending on the parameter values. An interesting but technically challenging direction for future work would be to make systematic modifications of natural image patches, for instance, enhancing texture differences or blurring the boundary, to explore the effects this has on edge category classification. A somewhat crude version of natural image patch manipulation in the form of removing texture information entirely has been performed in our previous work to establish the importance of texture as a cue for edge detection and classification (**DiMattina et al., 2012, 2022**).

Another limitation to this work is that we did not attempt here to establish the relative importance of various cues for classification. Doing so properly would require a much more difficult experiment in which we manipulate each parameter in units of perceptual discriminability, as is often done in studies of cue combination (**Kingdom et al., 2015; Saarela & Landy, 2012; Hillis et al., 2004; Green & Swets, 1988**). The purpose of the present study was to simply establish in broad strokes that these parameters are important for human performance on the task, and it would be an interesting future study to better explore the relative importance of these parameters for classification. A large number of mathematical cue-combination models exist in the psychophysical and neuroscientific literature and could readily be applied to the current task (**Jones, 2016; Prins & Kingdom, 2018**) in a refinement of the current study.

Additional limitations of any experiment seeking to understand the contributions that local cues make to perceptual organization, is that local stimulus processing is profoundly influenced by global context (**Schwartz, Hsu, & Dayan, 2007**; **Neri, 2011; Gilbert & Li, 2013; Spillman, Dresp-Langley, & Tseng 2015**). For this reason, it is likely that our study under-estimates classification performance, since in natural perception the local image would not be presented in isolation but with contextual information as well. However, the utility of our approach is that it demonstrates the sufficiency of locally available cues for image parsing, providing hints to workers in neurophysiology where in the visual processing hierarchy one may find neurons that first contribute information for image parsing, even if the task is ultimately performed upstream.

Finally, in our opinion the most interesting direction for future research lies not in psychophysics or computational studies, but in visual neurophysiology. One of the essential computations that the visual system performs is discounting the illuminant, enabling accurate identification of surfaces and their boundary without regard to shading or cast shadows (**Kingdom, 2003, 2008; Murray 2021**). However, it is unknown exactly where in the visual system this discounting of the illuminant takes place. Our work here and elsewhere (**DiMattina et al., 2022**) demonstrates that information useful to this process may well take place relatively early in the processing hierarchy, as small image patches (40 × 40 pixels) subtending only a few degrees of visual angle seem to be reliably classified as changes in material or illumination. It would be of great interest for future work in neurophysiology to determine if neurons in these areas responsive to our stimulus set exhibit the same ability to reliably distinguish image the categories considered here (occlusion, shadow, texture) via population decoding of these visual regions. A hypothetical neurophysiology experiment which directly complements with the psychophysics here would be to present to a population of neurons all of the stimuli from the two training sets (**TRN-1**, **TRN-2**), and then train a machine-learning classifier to decode the stimulus category from the neural population responses. Then, present the neural population with stimuli from the test-sets (**TST-1**,…,**TST-5**) and use the elicited neural responses as inputs to the trained classifier to obtain category predictions which can be directly compared with our psycho-physical data. A strong agreement between decoded neural responses and visual behavior would provide a strong argument that the neural population in question may underlie the ability of the visual system to perform this essential perceptual task. Given the fact that neurons in visual area V4 are sensitive to form (**Pasupathy & Connor, 2002**), texture (**Kim, Bair, & Pasupathy, 2019, 2022; Okazawa, Tajima, & Komatsu, 2017**) and blur (**Oleskiw, Nowack, & Pasupathy, 2018**), this may be a reasonable place to look for neurons that can reliably detect edges and accurately classify them as arising from shadows or occlusions.

## Acknowledgements

We thank FGCU undergraduates in the Computational Perception Lab for help with data collection.

## REFERENCES

Akaike, H. (1974). A new look at the statistical model identification. IEEE Transactions on Automatic Control, 19(6), 716–723.

Bishop, C. M., & Nasrabadi, N. M. (2006). Pattern recognition and machine learning (Vol. 4, No. 4, p. 738). New York: Springer.

Boussaoud, D., Desimone, R., & Ungerleider, L. G. (1991). Visual topography of area TEO in the macaque. Journal of Comparative Neurology, 306(4), 554–575.

Breuil, C., Jennings, B. J., Barthelmé, S., Guyader, N., & Kingdom, F. A. (2019). Color improves edge classification in human vision. PLoS Computational Biology, 15(10), e1007398.

Burge, J., Fowlkes, C. C., & Banks, M. S. (2010). Natural-scene statistics predict how the figure– ground cue of convexity affects human depth perception. Journal of Neuroscience, 30(21), 7269–7280.

DiMattina, C., Burnham, J. J., Guner, B. N., & Yerxa, H. B. (2022). Distinguishing shadows from surface boundaries using local achromatic cues. PLoS Computational Biology, 18(9), e1010473.

DiMattina, C., Fox, S. A., & Lewicki, M. S. (2012). Detecting natural occlusion boundaries using local cues. Journal of Vision, 12(13), 15–15.

DiMattina, C., & Wang, X. (2006). Virtual vocalization stimuli for investigating neural representations of species-specific vocalizations. Journal of Neurophysiology, 95(2), 1244–1262.

Elder, J. H., & Zucker, S. W. (1998). Local scale control for edge detection and blur estimation. IEEE Transactions on Pattern Analysis and Machine Intelligence, 20(7), 699–716.

Elston, G. N., & Rosa, M. G. (1998). Morphological variation of layer III pyramidal neurones in the occipitotemporal pathway of the macaque monkey visual cortex. Cerebral Cortex (New York, NY: 1991), 8(3), 278-294.

Ehinger, K. A., Adams, W. J., Graf, E. W., & Elder, J. H. (2017). Local depth edge detection in humans and deep neural networks. In Proceedings of the IEEE International Conference on Computer Vision Workshops (pp. 2681-2689).

Fowlkes, C. C., Martin, D. R., & Malik, J. (2007). Local figure–ground cues are valid for natural images. Journal of Vision, 7(8), 2–2.

Frisby, J. P., & Stone, J. V. (2010). Seeing: The Computational Approach to Biological Vision. MIT Press.

Gilbert, C. D., & Li, W. (2013). Top-down influences on visual processing. Nature Reviews Neuroscience, 14(5), 350–363.

Green, D. M., & Swets, J. A. (1966). Signal Detection Theory and Psychophysics (Vol. 1, pp. 1969-2012). New York: Wiley.

Hansen, T., & Gegenfurtner, K. R. (2009). Independence of color and luminance edges in natural scenes. Visual Neuroscience, 26(1), 35–49.

Hansen, T., & Gegenfurtner, K. R. (2017). Color contributes to object-contour perception in natural scenes. Journal of Vision, 17(3), 14–14.

Hillis, J. M., Watt, S. J., Landy, M. S., & Banks, M. S. (2004). Slant from texture and disparity cues: Optimal cue combination. Journal of Vision, 4(12), 1–1.

Ing, A. D., Wilson, J. A., & Geisler, W. S. (2010). Region grouping in natural foliage scenes: Image statistics and human performance. Journal of Vision, 10(4), 10–10.

Jing, J., Liu, S., Wang, G., Zhang, W., & Sun, C. (2022). Recent advances on image edge detection: A comprehensive review. Neurocomputing, 503, 259–271.

Jones, P. R. (2016). A tutorial on cue combination and Signal Detection Theory: Using changes in sensitivity to evaluate how observers integrate sensory information. Journal of Mathematical Psychology, 73, 117–139.

Kim, T., Bair, W., & Pasupathy, A. (2019). Neural coding for shape and texture in macaque area V4. Journal of Neuroscience, 39(24), 4760–4774.

Kim, T., Bair, W., & Pasupathy, A. (2022). Perceptual texture dimensions modulate neuronal response dynamics in visual cortical area V4. Journal of Neuroscience, 42(4), 631–642.

Kingdom, F. A. (2003). Color brings relief to human vision. Nature Neuroscience, 6(6), 641–644.

Kingdom, F. A. (2008). Perceiving light versus material. Vision Research, 48(20), 2090-2105.

Kobatake, E., & Tanaka, K. (1994). Neuronal selectivities to complex object features in the ventral visual pathway of the macaque cerebral cortex. Journal of Neurophysiology, 71(3), 856–867.

Konishi, S., Yuille, A. L., Coughlan, J. M., & Zhu, S. C. (2003). Statistical edge detection: Learning and evaluating edge cues. IEEE Transactions on Pattern Analysis and Machine Intelligence, 25(1), 57–74.

Liberman, A. M. (1996). Speech: A special code. MIT press.

Loffler, G., Yourganov, G., Wilkinson, F., & Wilson, H. R. (2005). fMRI evidence for the neural representation of faces. Nature Neuroscience, 8(10), 1386–1391.

Margoliash, D., & Fortune, E. S. (1992). Temporal and harmonic combination-sensitive neurons in the zebra finch’s HVc. Journal of Neuroscience, 12(11), 4309–4326.

Marr, D. (1982). Vision: A computational investigation into the human representation and processing of visual information. MIT press.

Martin, D. R., Fowlkes, C. C., & Malik, J. (2004). Learning to detect natural image boundaries using local brightness, color, and texture cues. IEEE Transactions on Pattern Analysis and Machine Intelligence, 26(5), 530–549.

Mély, D. A., Kim, J., McGill, M., Guo, Y., & Serre, T. (2016). A systematic comparison between visual cues for boundary detection. Vision Research, 120, 93–107.

Murphy, K. P. (2012). Machine learning: a probabilistic perspective. MIT press.

Murray, R. F. (2021). Lightness perception in complex scenes. Annual Review of Vision Science, 7(1), 417–436.

Neri, P. (2011). Global properties of natural scenes shape local properties of human edge detectors. Frontiers in Psychology, 2, 172.

Okazawa, G., Tajima, S., & Komatsu, H. (2017). Gradual development of visual texture-selective properties between macaque areas V2 and V4. Cerebral Cortex, 27(10), 4867–4880.

Oleskiw, T. D., Nowack, A., & Pasupathy, A. (2018). Joint coding of shape and blur in area V4. Nature Communications, 9(1), 1–13.

Olmos, A., & Kingdom, F. A. (2004). A biologically inspired algorithm for the recovery of shading and reflectance images. Perception, 33(12), 1463–1473.

O’Neill, W. E., & Suga, N. (1979). Target range-sensitive neurons in the auditory cortex of the mustache bat. Science, 203(4375), 69–73.

Op De Beeck, H., & Vogels, R. (2000). Spatial sensitivity of macaque inferior temporal neurons. Journal of Comparative Neurology, 426(4), 505–518.

Osmanski, M. S., & Wang, X. (2023). Perceptual specializations for processing species-specific vocalizations in the common marmoset (Callithrix jacchus). Proceedings of the National Academy of Sciences, 120(24), e2221756120.

Pasupathy, A., & Connor, C. E. (2002). Population coding of shape in area V4. Nature Neuroscience, 5(12), 1332–1338.

Peterson, J. C., Uddenberg, S., Griffiths, T. L., Todorov, A., & Suchow, J. W. (2022). Deep models of superficial face judgments. Proceedings of the National Academy of Sciences, 119(17), e2115228119.

Prins, N., & Kingdom, F. A. (2018). Applying the model-comparison approach to test specific research hypotheses in psychophysical research using the Palamedes toolbox. Frontiers in Psychology, 9, 1250.

Ramenahalli, S., Mihalas, S., & Niebur, E. (2014). Local spectral anisotropy is a valid cue for figure–ground organization in natural scenes. Vision Research, 103, 116–126.

Saarela, T., & Landy, M. (2012). Integration of texture and color cues for visual shape recognition. Journal of Vision, 12(9), 102–102.

Schwartz, O., Hsu, A., & Dayan, P. (2007). Space and time in visual context. Nature Reviews Neuroscience, 8(7), 522–535.

Spillmann, L., Dresp-Langley, B., & Tseng, C. H. (2015). Beyond the classical receptive field: The effect of contextual stimuli. Journal of Vision, 15(9), 7–7.

Vilankar, K. P., Golden, J. R., Chandler, D. M., & Field, D. J. (2014). Local edge statistics provide information regarding occlusion and nonocclusion edges in natural scenes. Journal of Vision, 14(9), 13–13.

Wilson, H. R., & Wilkinson, F. (2015). From orientations to objects: Configural processing in the ventral stream. Journal of Vision, 15(7), 4–4.

Zhou, C., & Mel, B. W. (2008). Cue combination and color edge detection in natural scenes. Journal of Vision, 8(4), 4–4.

