## Supplementary_Material for "Local cues enable classification of image patches as surfaces, object boundaries, or illumination changes"

**SUPPLEMENTARY FIGURE S1**

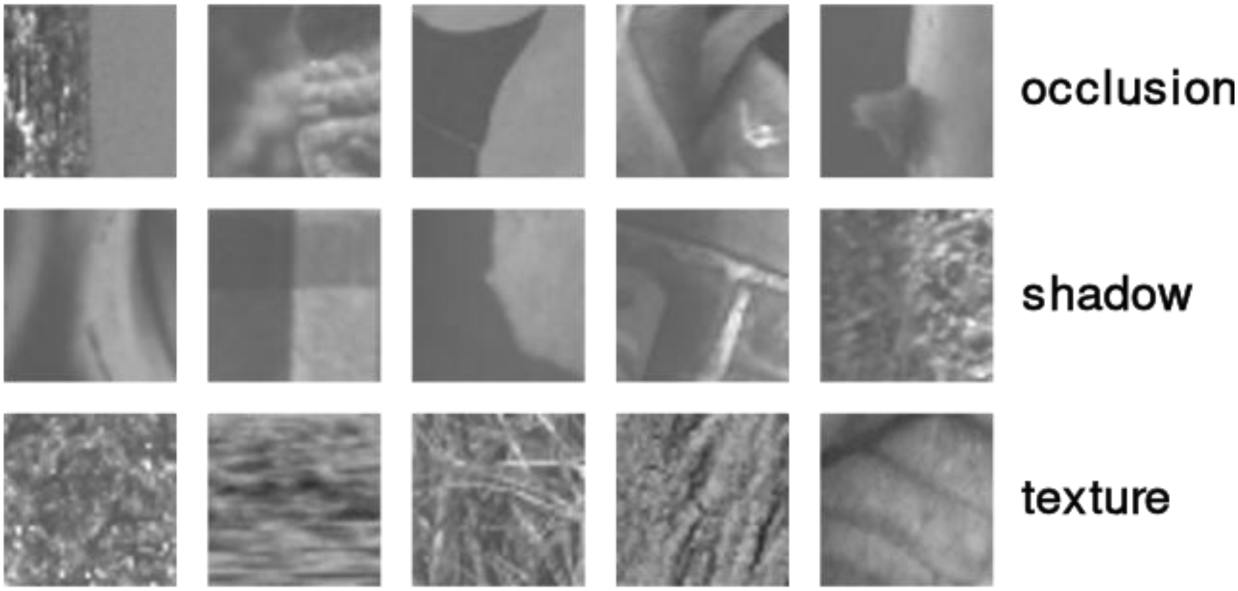


**Supplementary Fig. S1:** Examples of stimuli from Training Survey 2 (**TRN-2**) from each of the three categories.

### SUPPLEMENTARY FIGURE S2

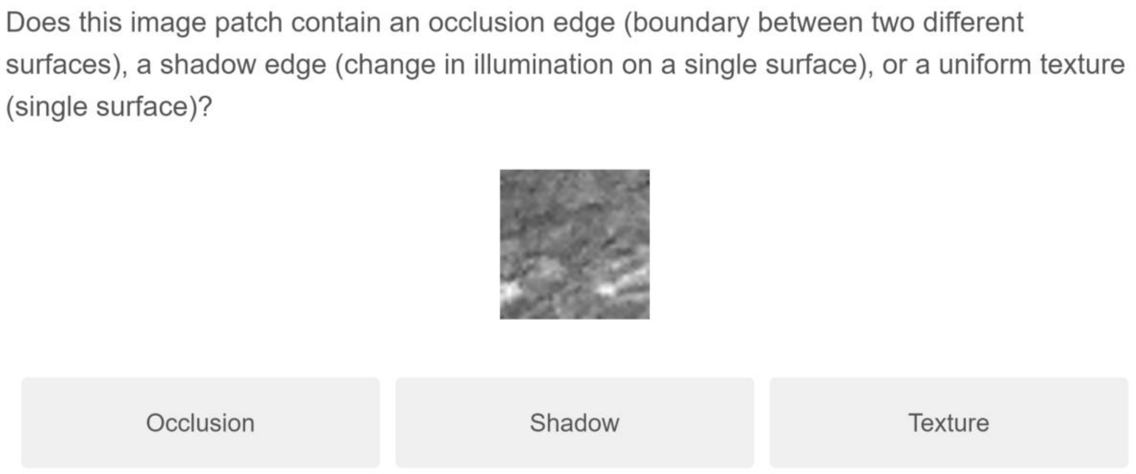


**Supplementary Fig. S2:** Sample question from Training Survey 1 (**TRN-1**).

### SUPPLEMENTARY FIGURE S3

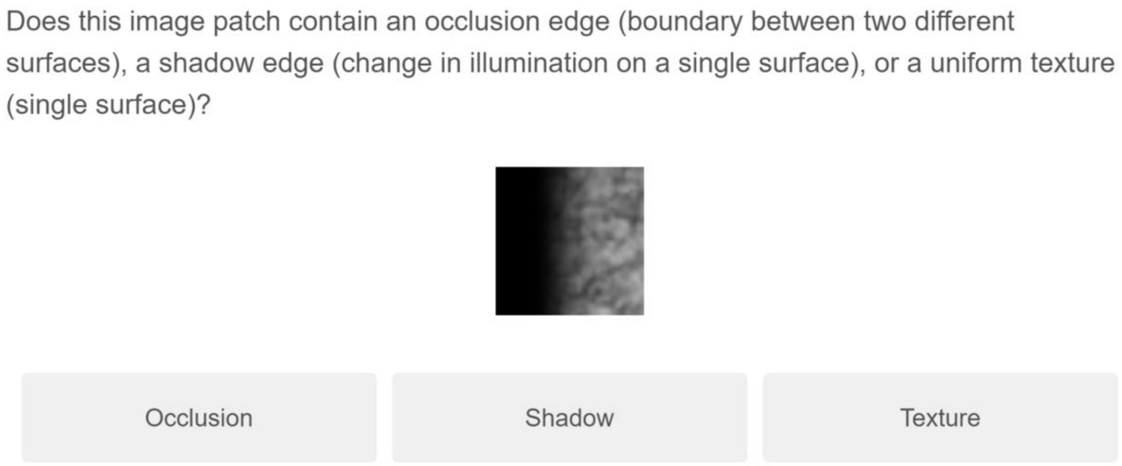


**Supplementary Fig. S3:** Sample question from Test Survey 5 (**TST-5**).

**SUPPLEMENTARY FIGURE S4**

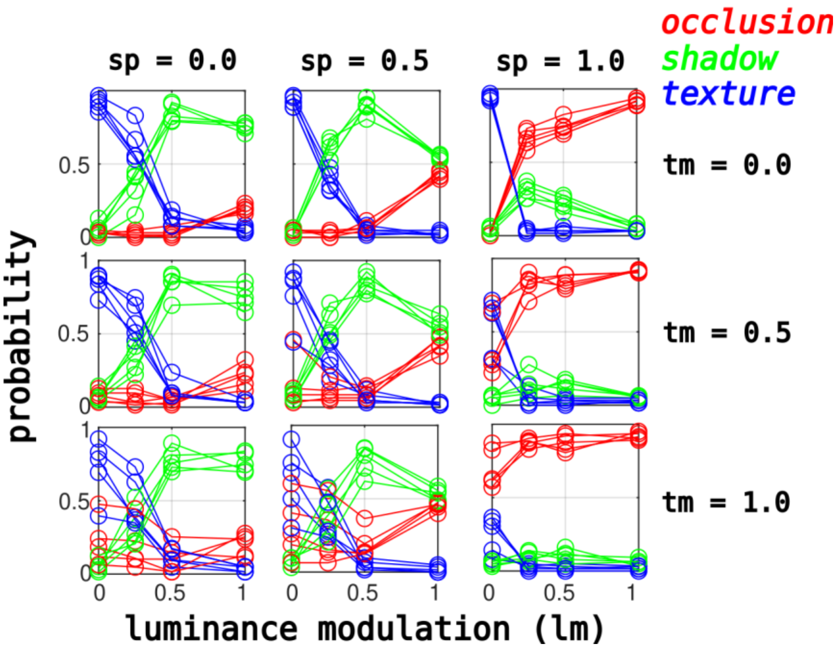


**Supplementary Fig. S4:** Same as **Fig. 5**, but for individual surveys.

**SUPPLEMENTARY FIGURE S5**

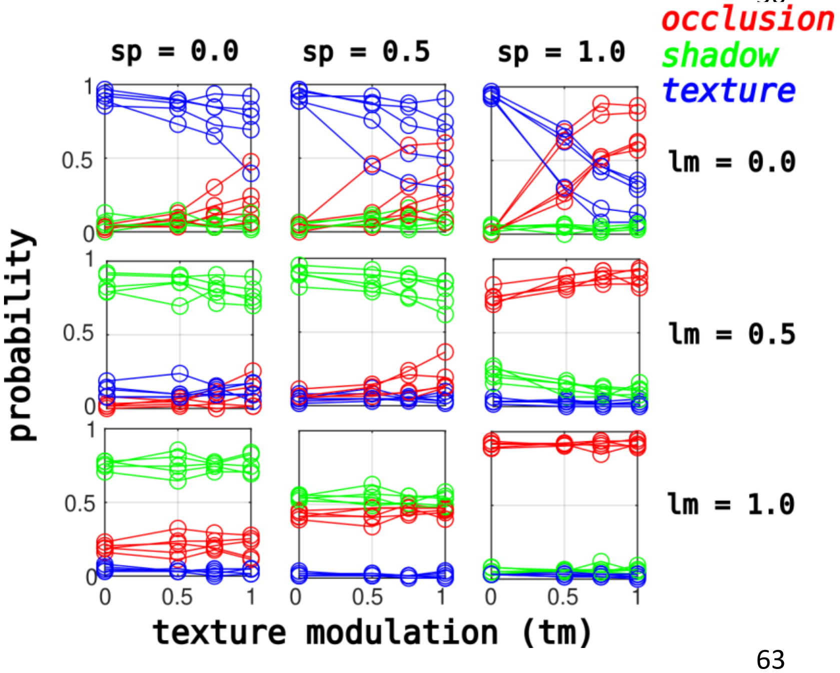


**Supplementary Fig. S5:** Same as **Fig. 6**, but for individual surveys.

### SUPPLEMENTARY FIGURE S6

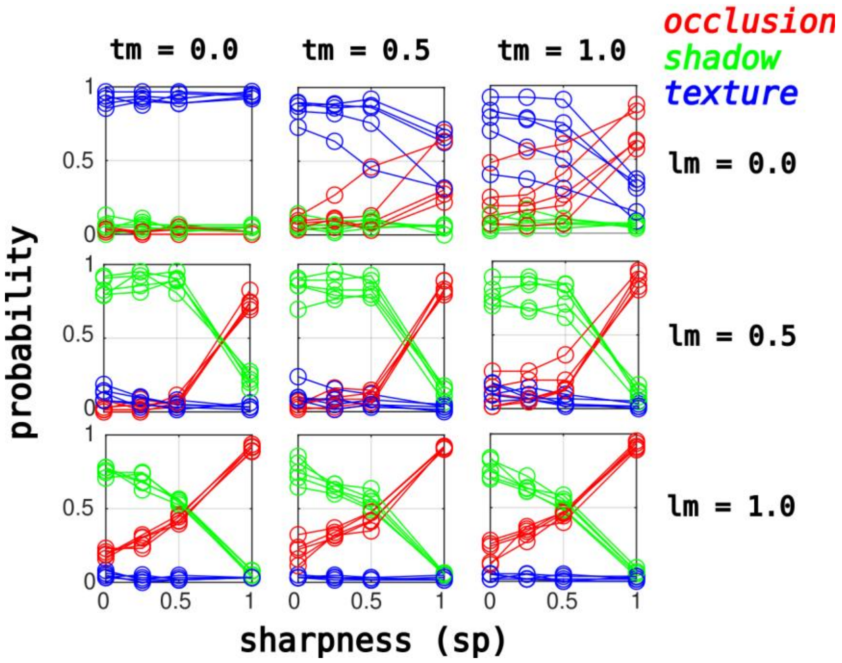


**Supplementary Fig. S6:** Same as **Fig. 7**, but for individual surveys.

**SUPPLEMENTARY FIGURE S7**

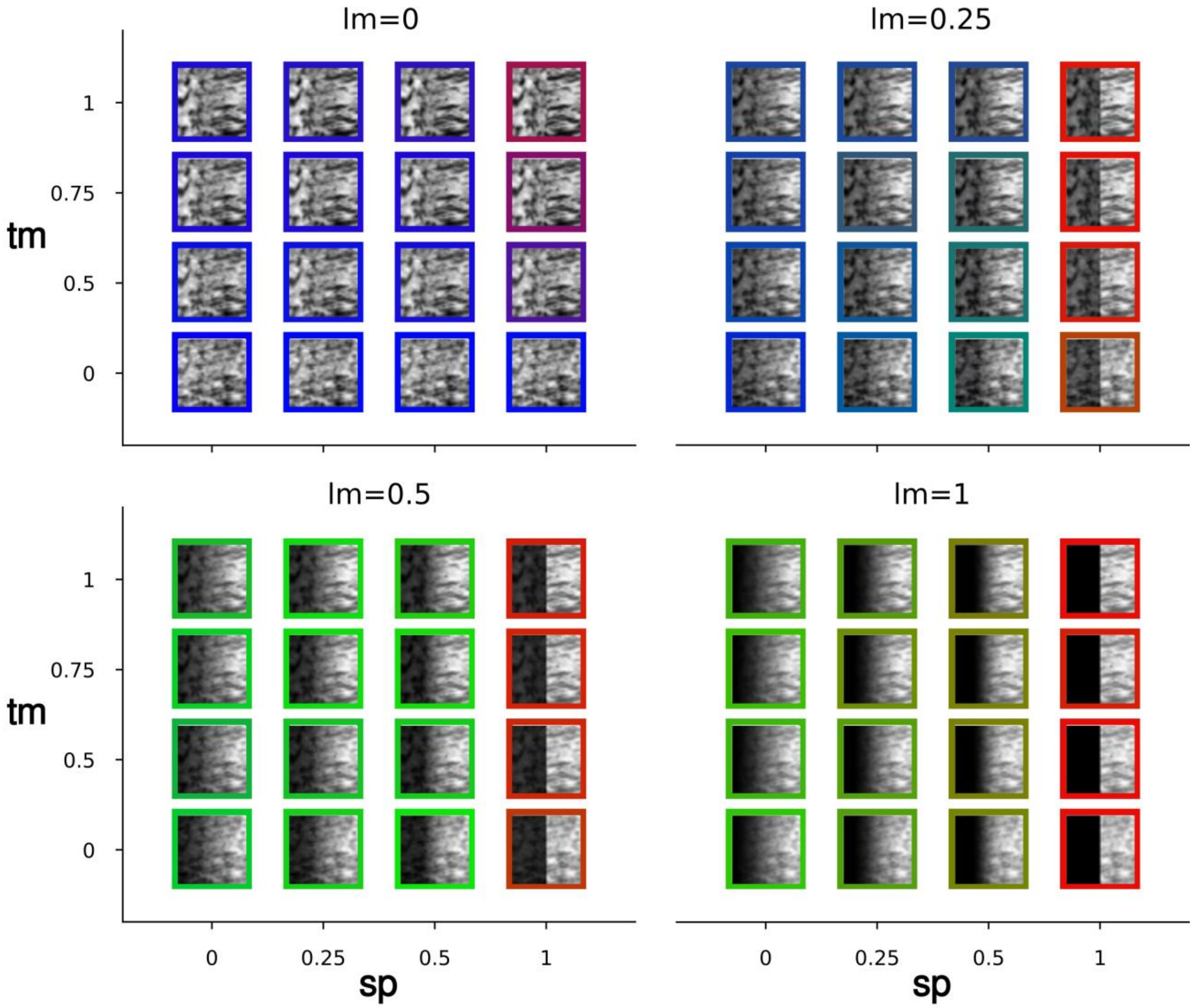


**Supplementary Fig. S7:** Same as **Fig. 8** but for **TST-2**.

**SUPPLEMENTARY FIGURE S8**

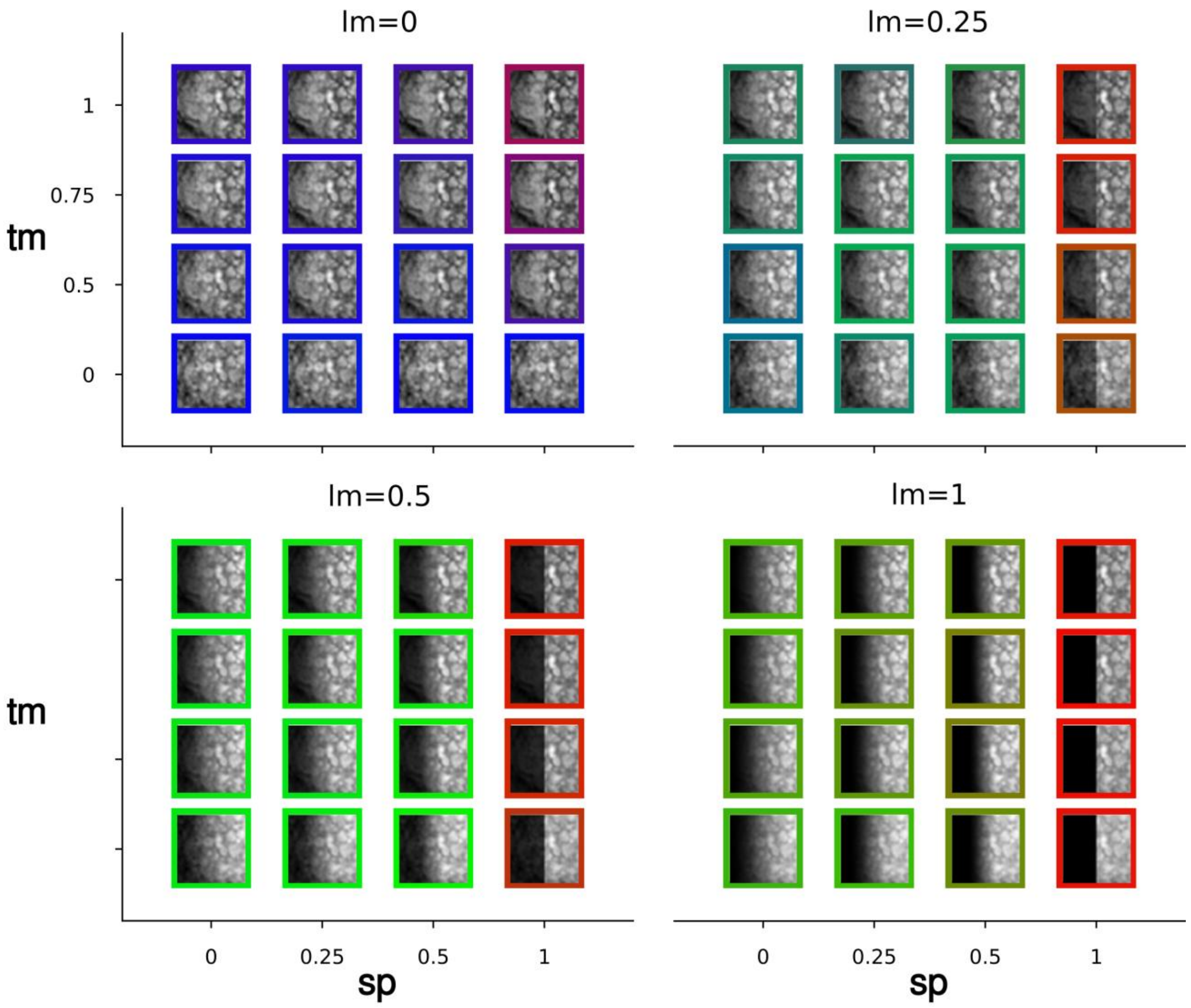


**Supplementary Fig. S8:** Same as **Fig. 8** but for **TST-3**.

**SUPPLEMENTARY FIGURE S9**

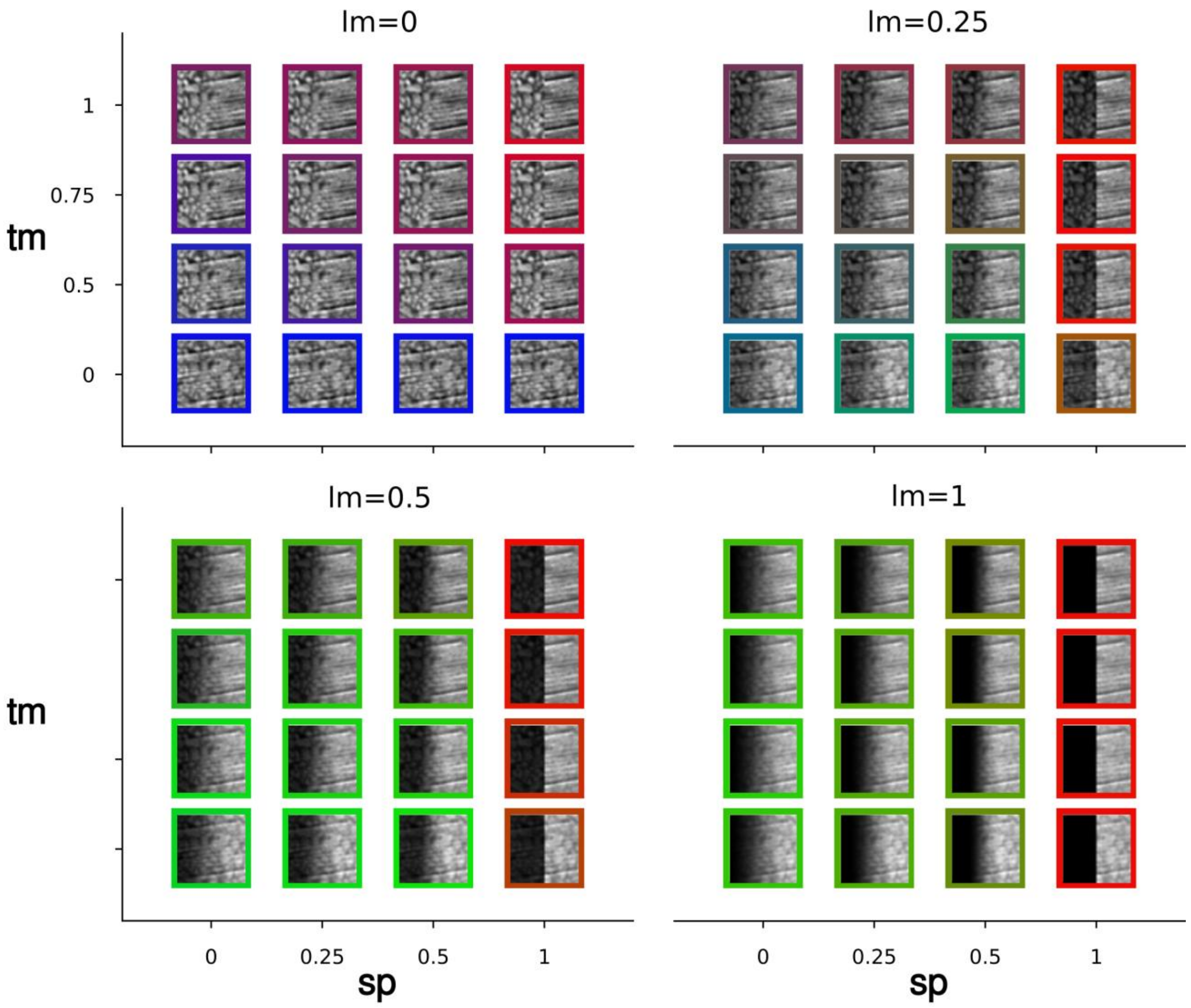


**Supplementary Fig. S9:** Same as **Fig. 8** but for **TST-4**.

**SUPPLEMENTARY FIGURE S10**

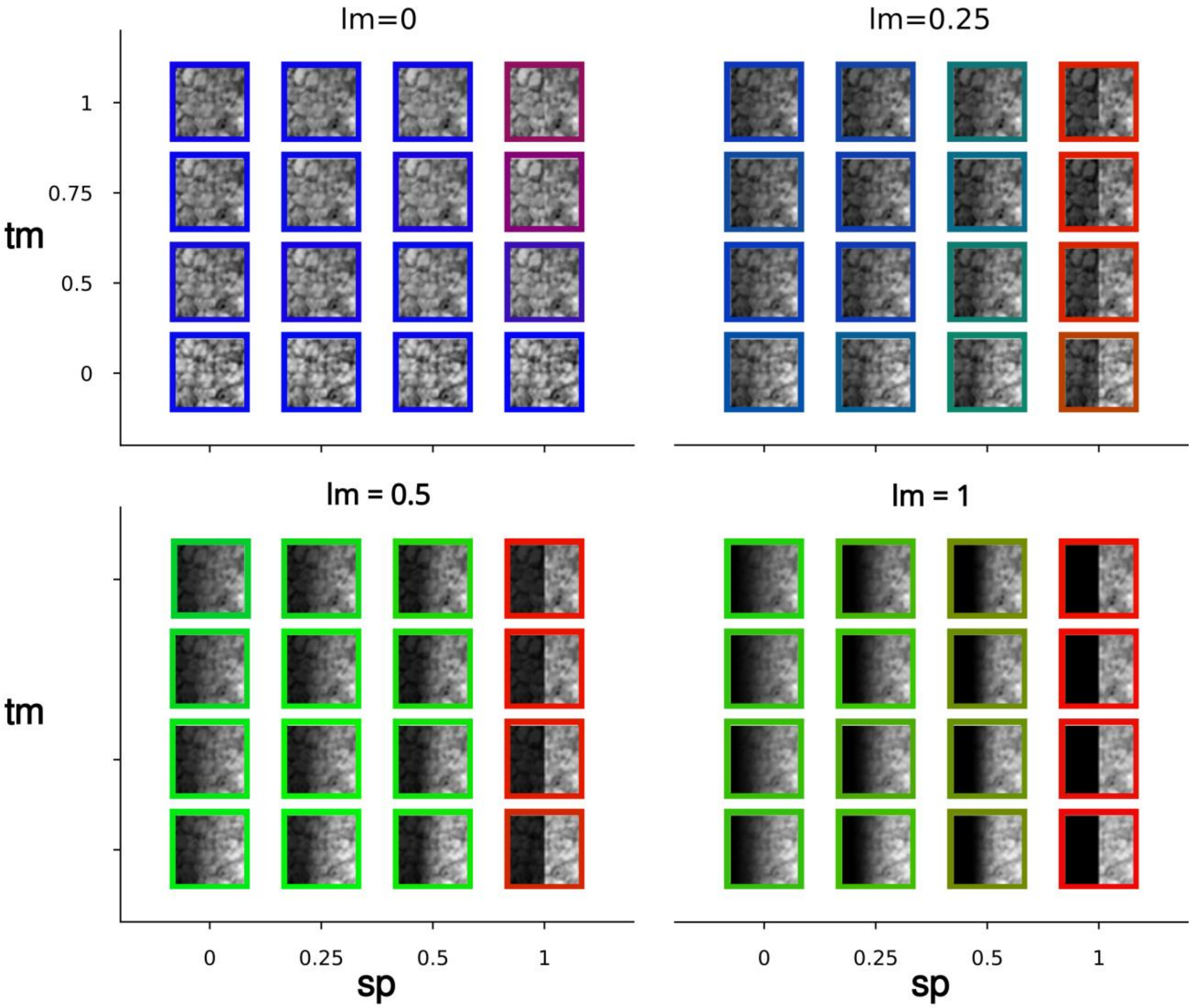


**Supplementary Fig. S10:** Same as **Fig. 8** but for **TST-5**.

**SUPPLEMENTARY FIGURE S11**

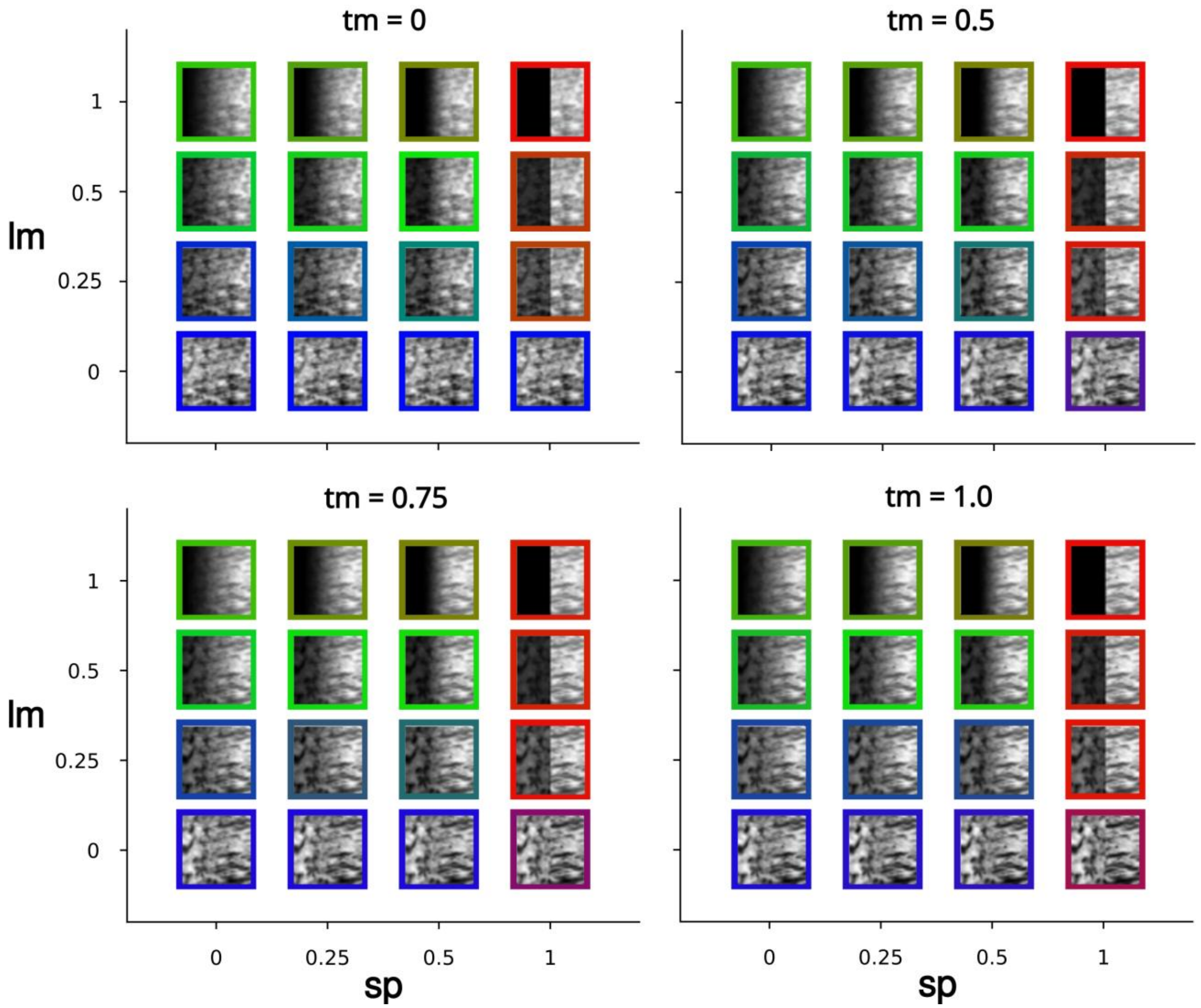


**Supplementary Fig. S11:** Same as **Fig. 9** but for **TST-2**.

**SUPPLEMENTARY FIGURE S12**

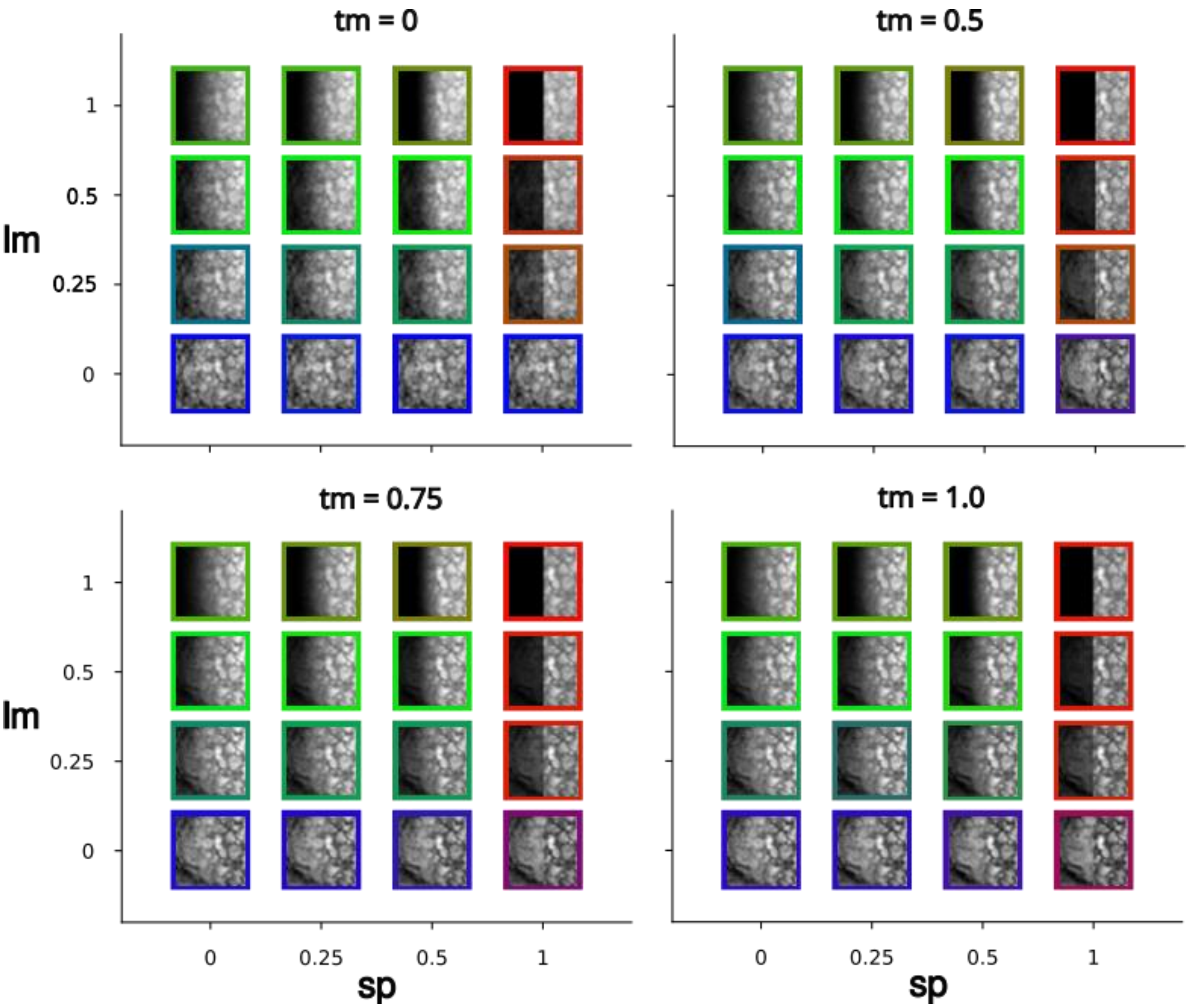


**Supplementary Fig. S12:** Same as **Fig. 9** but for **TST-3**.

**SUPPLEMENTARY FIGURE S13**

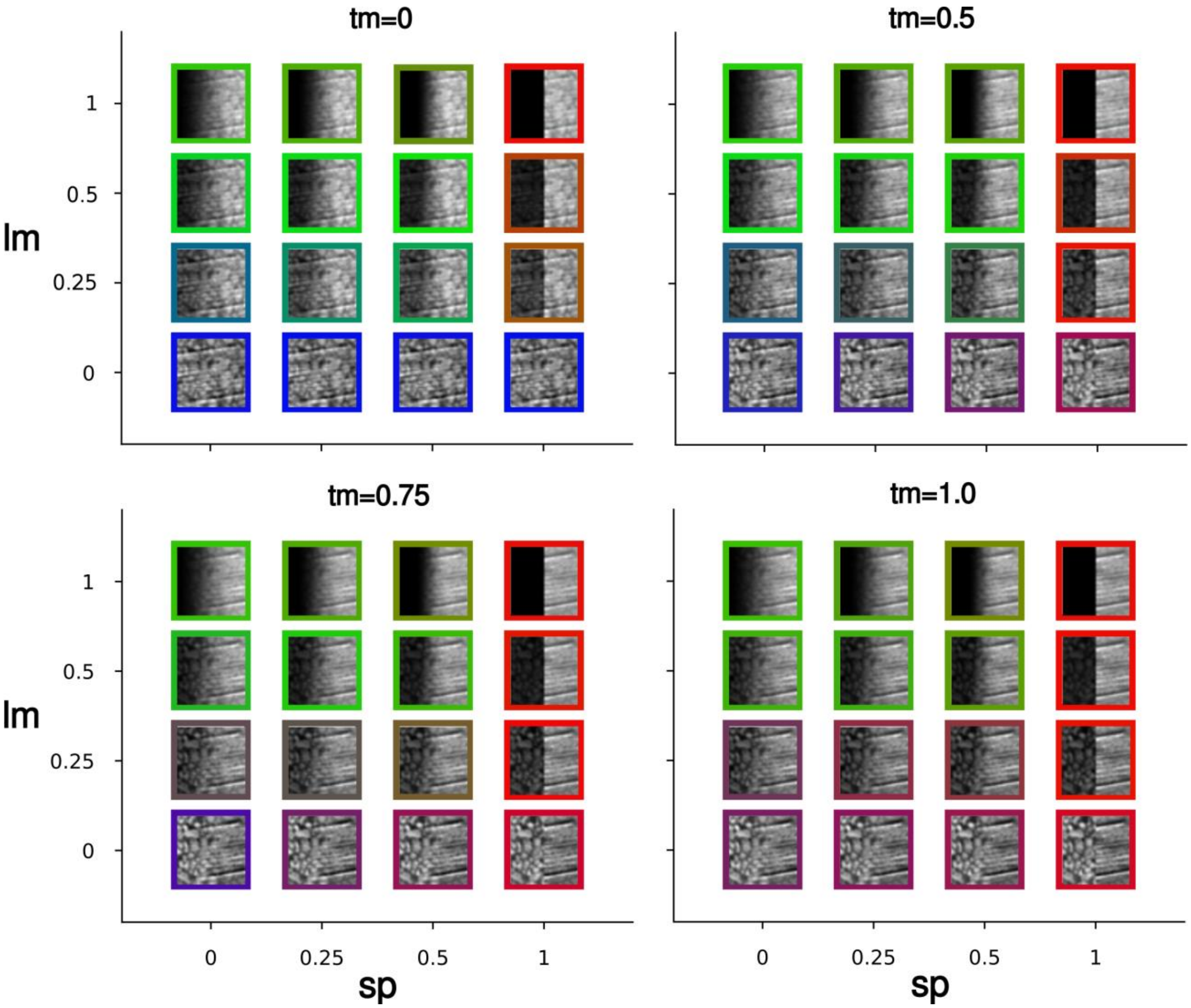


**Supplementary Fig. S13:** Same as **Fig. 9** but for **TST-4**.

**SUPPLEMENTARY FIGURE S14**

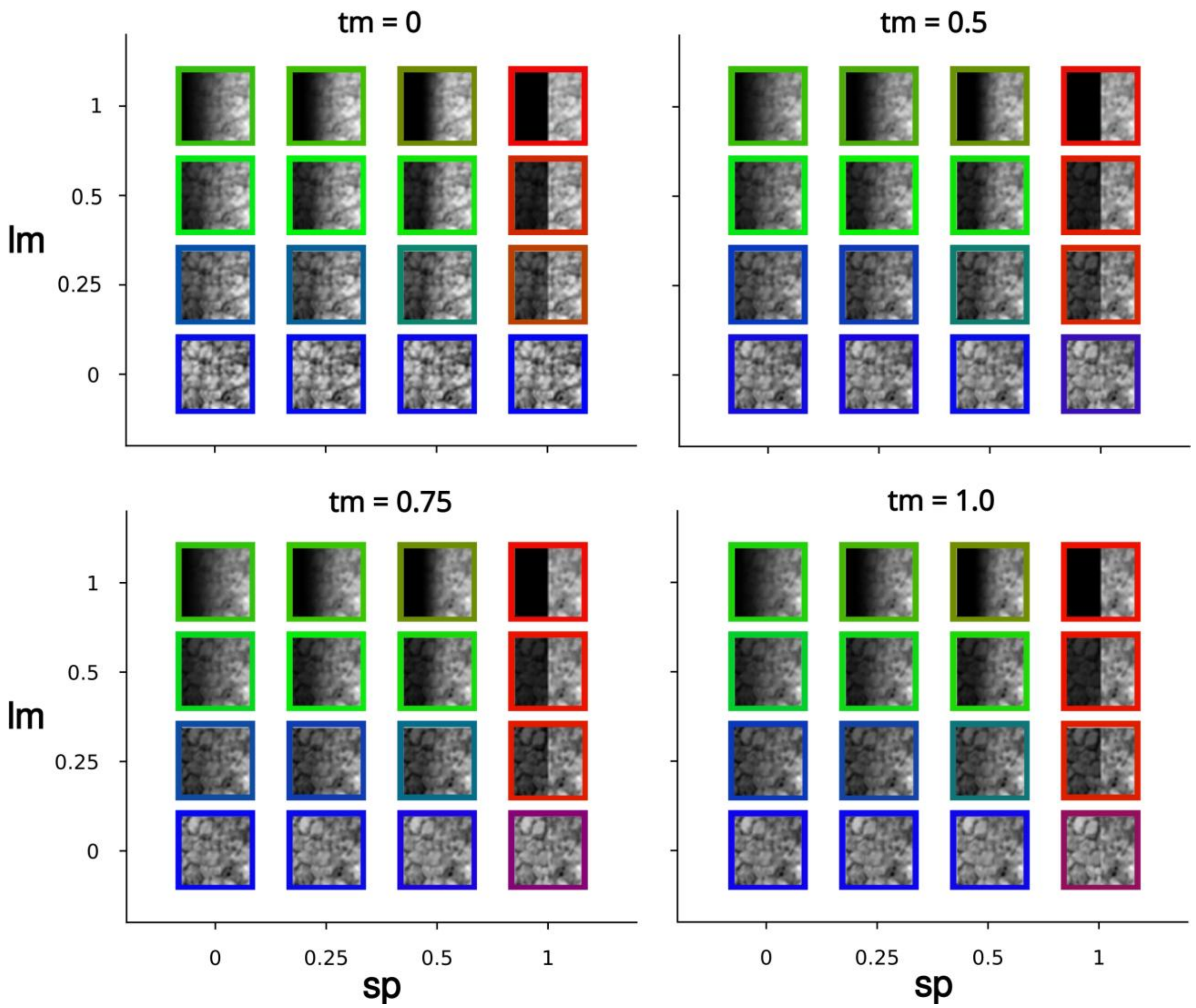


**Supplementary Fig. S14:** Same as **Fig. 9** but for **TST-5**.

**SUPPLEMENTARY FIGURE S15**

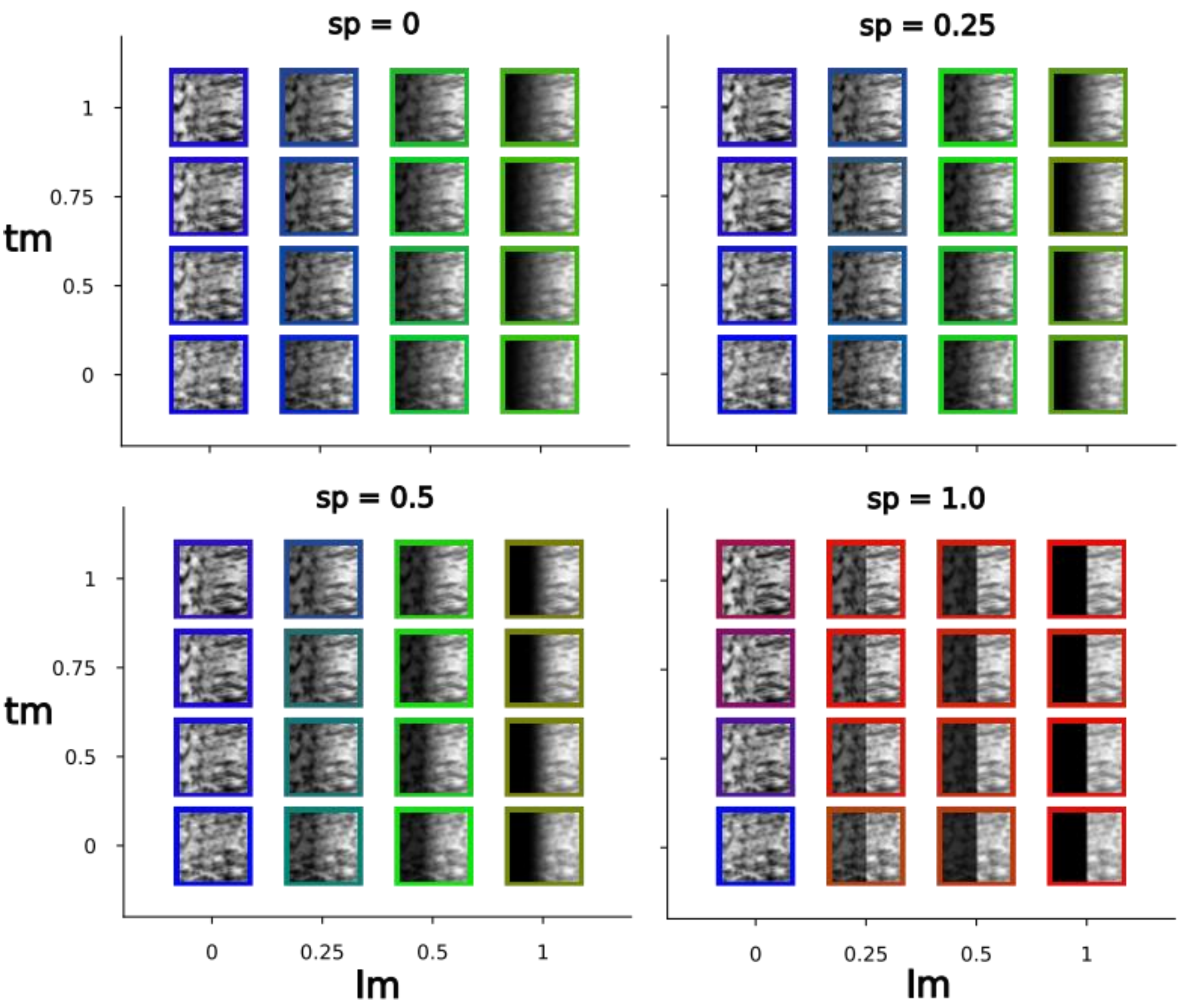


**Supplementary Fig. S15:** Same as **Fig. 10** but for **TST-2**.

**SUPPLEMENTARY FIGURE S16**

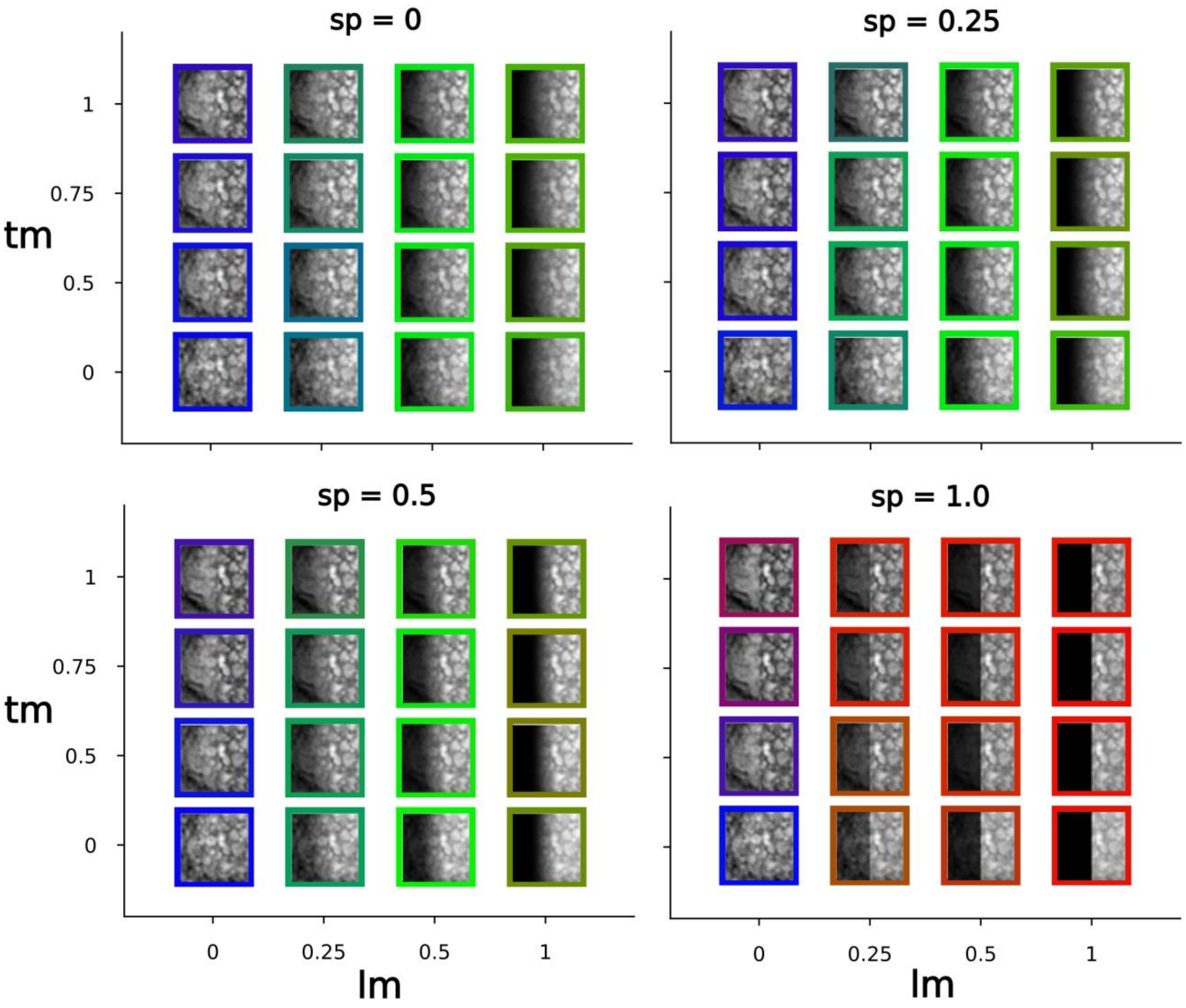


**Supplementary Fig. S16:** Same as **Fig. 10** but for **TST-3**.

**SUPPLEMENTARY FIGURE S17**

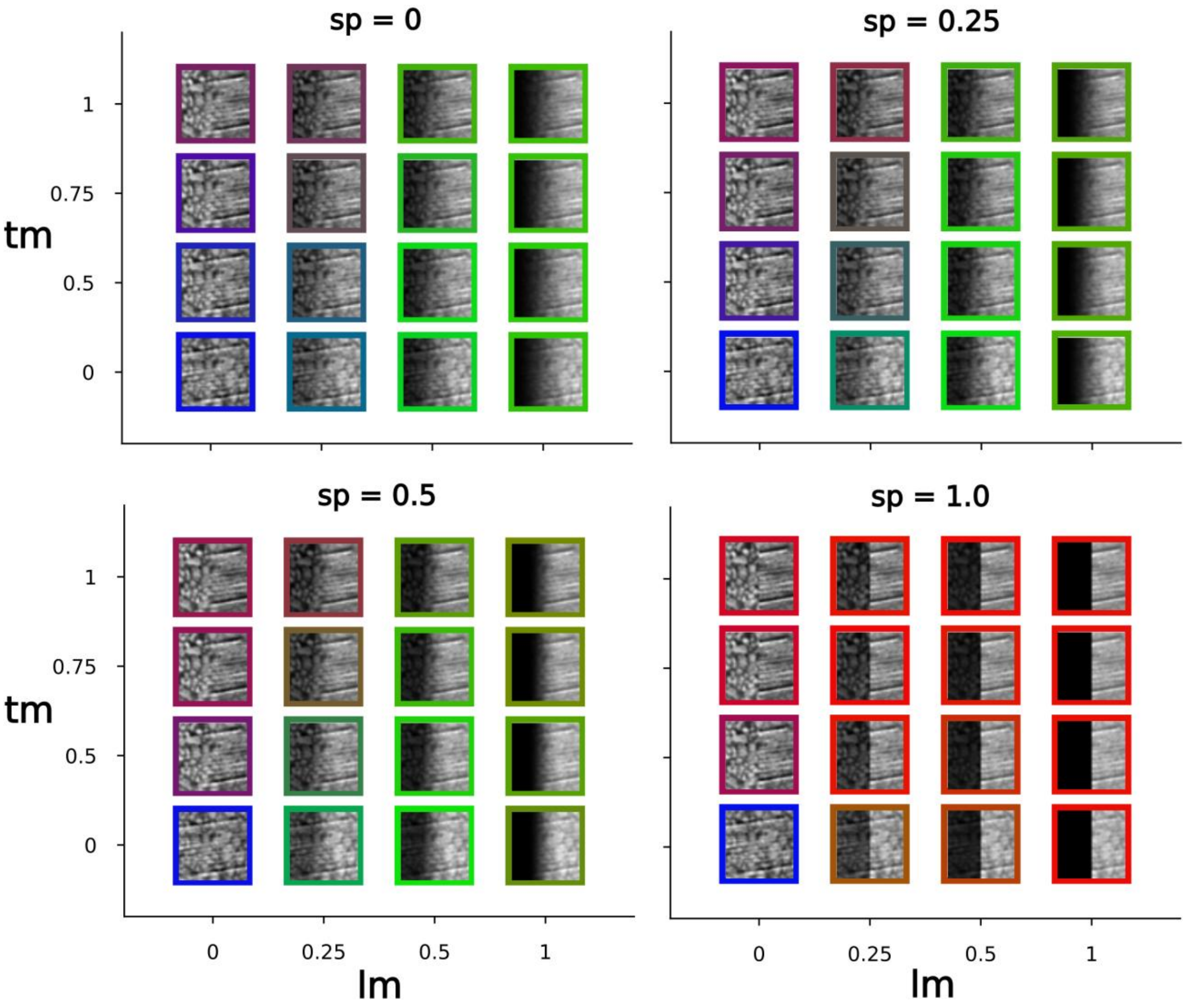


**Supplementary Fig. S17**: Same as **Fig. 10** but for **TST-4**.

**SUPPLEMENTARY FIGURE S18**

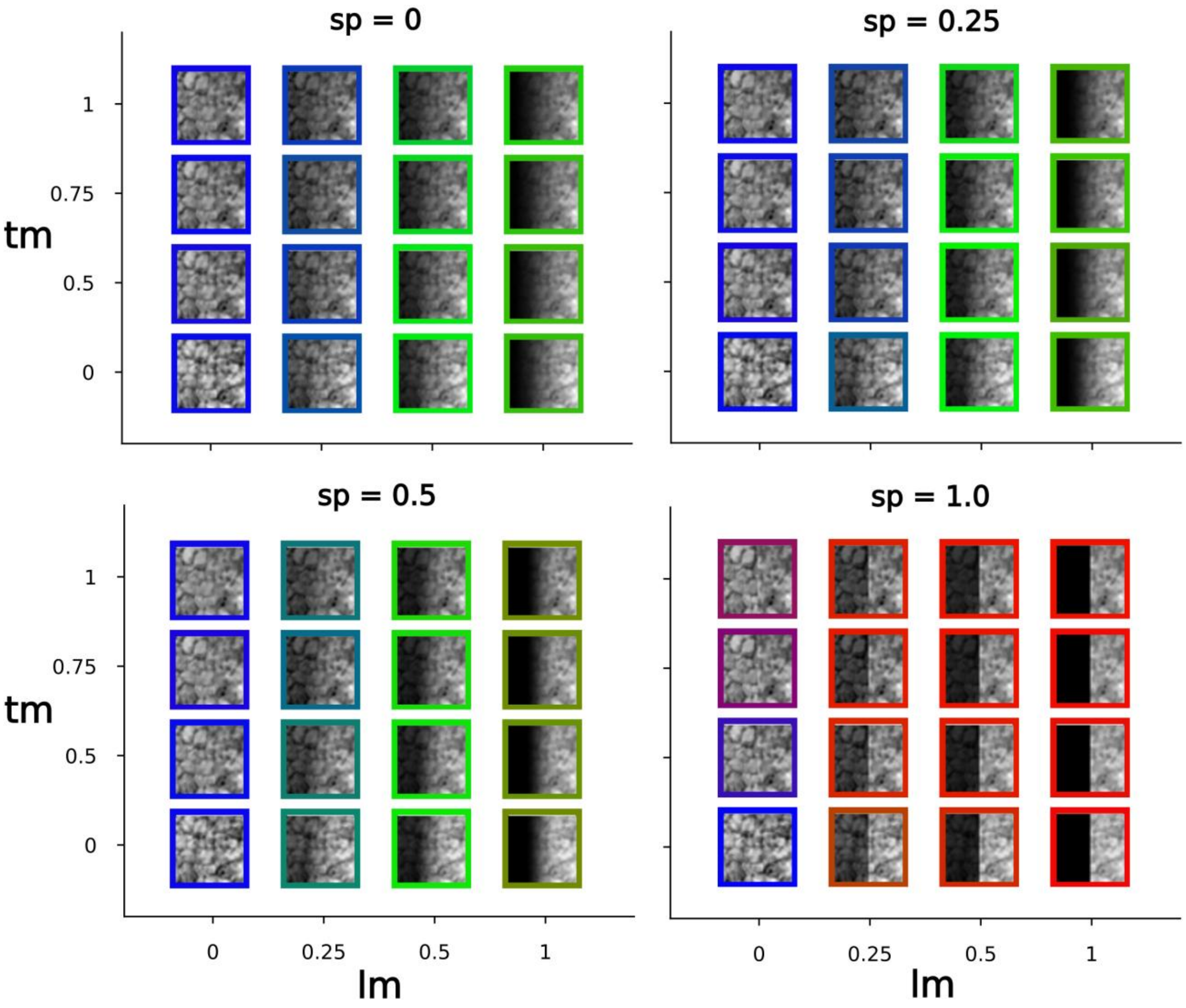


**Supplementary Fig. S18**: Same as **Fig. 10** but for **TST-5**.

**SUPPLEMENTARY FIGURE S19**

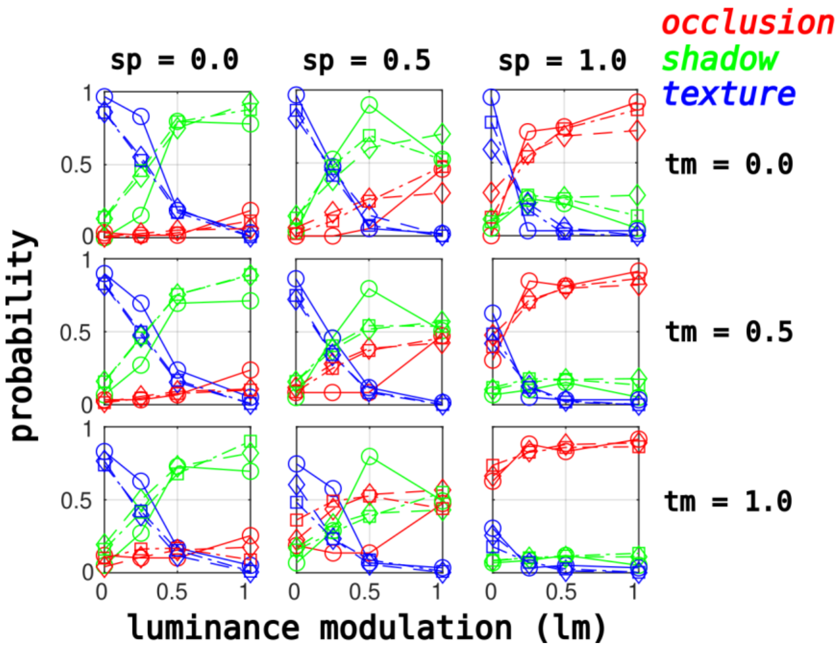


**Supplementary Fig. S19:** Same as **Fig. 11** but for **TST-2**.

**SUPPLEMENTARY FIGURE S20**

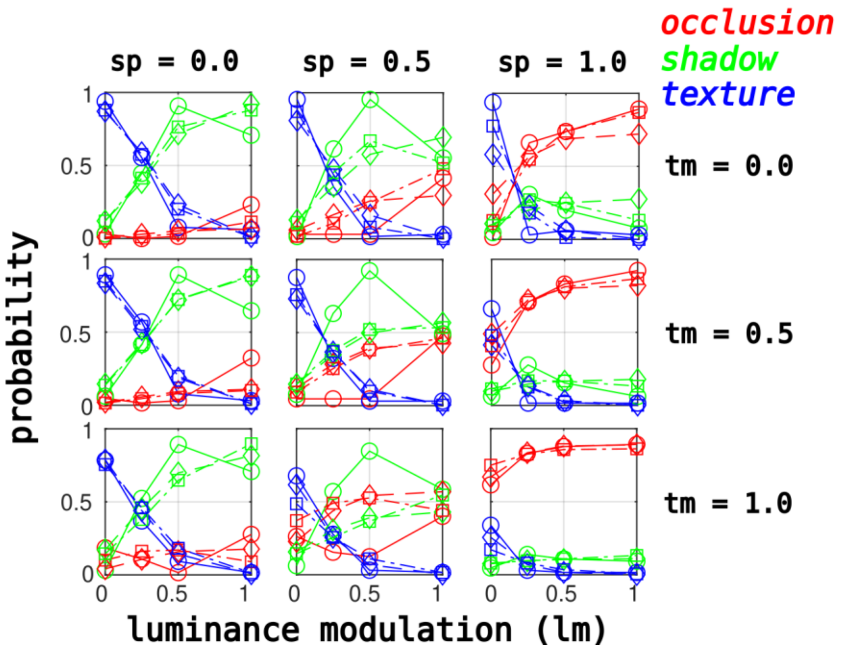


**Supplementary Fig. S20:** Same as **Fig. 11** but for **TST-3**.

**SUPPLEMENTARY FIGURE S21**

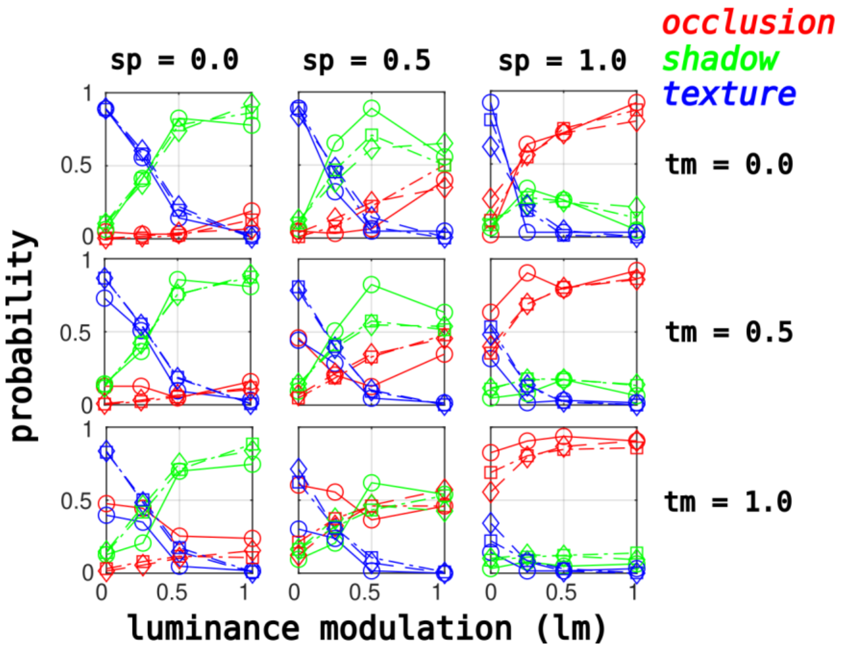


**Supplementary Fig. S21**: Same as **Fig. 11** but for **TST-4**.

**SUPPLEMENTARY FIGURE S22**

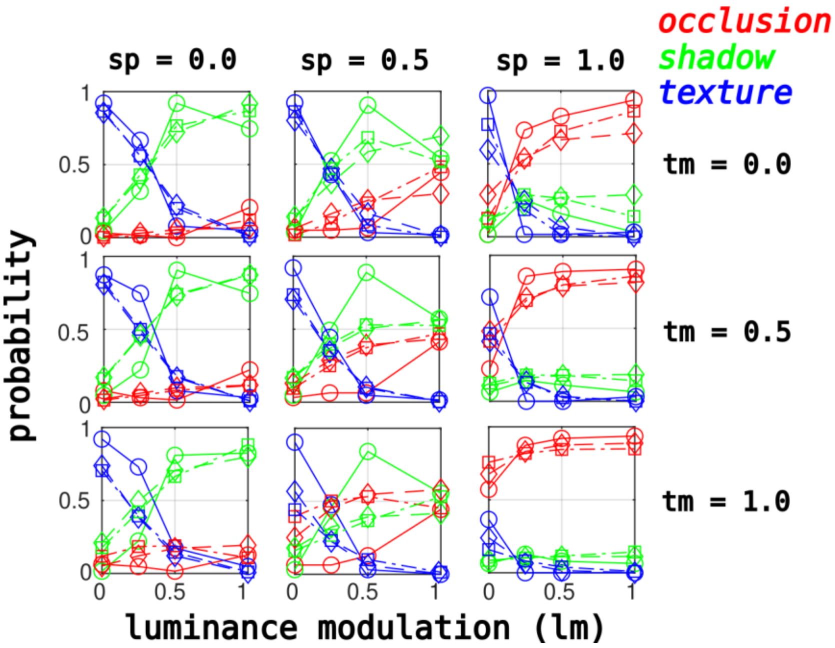


**Supplementary Fig. S22:** Same as **Fig. 11** but for **TST-5**.
